# Single-cell RNA sequencing of adult *Drosophila* ovary identifies transcriptional programs governing oogenesis

**DOI:** 10.1101/802314

**Authors:** Allison Jevitt, Deeptiman Chatterjee, Gengqiang Xie, Xian-Feng Wang, Taylor Otwell, Yi-Chun Huang, Wu-Min Deng

**Affiliations:** Department of Biological Science, Florida State University, Tallahassee, FL, 32306, USA; Department of Biochemistry and Molecular Biology, Tulane University School of Medicine, New Orleans, LA, 70112, USA

## Abstract

Oogenesis is a complex developmental process that involves spatiotemporally regulated coordination between the germline and supporting, somatic cell populations. This process has been modelled extensively using the *Drosophila* ovary. While different ovarian cell types have been identified through traditional means, the large-scale expression profiles underlying each cell type remain unknown. Using single-cell RNA sequencing technology, we have built a transcriptomic dataset for the adult *Drosophila* ovary and connected tissues. This dataset captures the entire transcriptional trajectory of the developing follicle cell population over time. Our findings provide detailed insight into processes such as cell-cycle switching, migration, symmetry breaking, nurse cell engulfment, egg-shell formation, and signaling during corpus luteum formation, marking a newly identified oogenesis-to-ovulation transition. Altogether, these findings provide a broad perspective on oogenesis at a single-cell resolution while revealing new genetic markers and fate-specific transcriptional signatures to facilitate future studies.

## Introduction

The adult *Drosophila* ovary is a versatile model used many biological studies. With powerful genetic tools available in *Drosophila*, studies of oogenesis have provided mechanistic insight into topics such as stem cell niche regulation [31, 60, 71, 76, 101], cell differentiation [3, 52], cell cycle and size control [9, 17], epithelial morphogenesis [46, 92, 100], cell migration [45, 66], tissue repair and homeostasis [88], etc. The success of this system as a developmental model is also due to the structure of the fly ovary, where eggs progress in sequence and many rounds of oogenesis occur simultaneously. This provides a unique advantage over other systems where temporal resolution and replicative power can be achieved easily within a single ovary.

A female fly has a pair of ovaries that are connected to the oviduct and held together by muscles known as the peritoneal sheath. Each ovary is made up of developmental units called ovarioles, which are individually sheathed within the musculature known as the epithelial sheath. Oogenesis occurs simultaneously within each of the 16-18 ovarioles, starting from stem cells at the anterior tip to the fully-developed eggs at the posterior end. Throughout oogenesis, the developing egg is supported by the germline-derived nurse cells, and the somatic follicular epithelium (made up of follicle cells). Together, the germline and the follicle cells form individual units called egg chambers. Egg chamber development is subdivided into early (1-6), middle (7-10A), and late (10B-14) stages based on mitotic, endocycle, and gene amplification cell-cycle programs of the follicle cells, respectively [47]. During ovulation, mature eggs break free from the epithelium and pass into the uterus through the oviduct. The epithelial layer remains in the ovary, forming a similar structure found in mammals, known as the corpus luteum [23].

To better understand how oogenesis is regulated at the cellular level, we performed single-cell RNA sequencing (scRNA-seq) on these ovarian cell types and uncovered novel gene expression patterns throughout oogenesis. With a special focus on the follicle cell trajectory we also described the major transcriptomic programs underlying the early, middle, and late stages of oogenesis. We also report a newly identified transcriptional shift in late-staged follicle cells (termed pre-corpus luteum cells) which begin upregulating ovulation-related genes.

## Materials and methods

### Experimental Model

#### Fly lines used for ScRNA-seq

All fly stocks and crosses were maintained at room temperature (23°C) and fed a yeast based medium. To construct the scRNA-seq dataset, *w*^−^ flies (BL#3605) were used, a common genetic background used in many studies [48].

#### Fly lines used in experimental validation of cluster markers

We used a variety of publicly available lines from Bloomington Stock Center to experimentally validate expression patterns of select genes from the scRNA-seq dataset. These lines fall into two categories: those with fluorescently tagged proteins under the control of a native promoter (either MiMIC-based RMCE [96] or protein trap [13]) and those expressing T2A-Gal4 (carrying either CRISPR-mediated insertions of T2A-Gal4 [57] or RMCE-mediated swap-ins of T2A-Gal4 [27]) driving UAS-GFP (BL#4775) or UAS-RFP (BL#31417) as a marker.

The GFP-tagged lines used in this study are Atf3:GFP (BL#42263), Ilp8:GFP (BL#33079), Past1:GFP (BL#51521), Glut4EF:GFP (BL#60555), abd-A:GFP (BL#68187), Chrac-16:GFP (BL#56160), shep:GFP (BL#61769), AdenoK:GFP (BL#56160), Fkbp1:GFP (BL#66358), mub:GFP (BL#51574), mnb:GFP (BL#66769), Gp210:GFP (BL#61651), Fpps:GFP (BL#51527), HmgD:GFP (BL#55827), sli:GFP (BL#64472), Nrx-IV:GFP (BL#50798), CG14207:GFP (BL#60226), D1:GFP (BL#66454), jumu:GFP (BL#59764), hdc:GFP (BL#59762), sm:GFP (BL#59815), Men:GFP (BL#61754), Sap-r:GFP (BL#63201), GILT1:GFP (BL#51543), Cp1:GFP (BL#51555). The T2A-Gal4 lines used in this study are Ance-Gal4 (BL#76676), FER-Gal4 (BL#67448), wb-Gal4 (BL#76189), stx-Gal4 (BL#77769), vir-1-Gal4 (BL#65650).

We also used Diap1:GFP, a kind gift from Jin Jiang Lab [105].

### Immunofluorescence and imaging

Ovaries and associated tissue were dissected in PBS, fixed for 15 minutes in 4% formaldehyde, washed 3 times in PBT, and then stained with DAPI (Invitrogen, 1:1000) to label nuclei. Samples were then mounted on slides in an 80% glycerol mounting solution. All images were captured using the Zeiss LSM 800 confocal microscope and associated Zeiss microscope software (ZEN blue).

### ScRNA-seq sample preparation

#### Dissociation and filtration of single cells

To maximize sampling genetic diversity between individuals, we dissected ovarian tissue from 50 adult flies. It is technically challenging to separate the ovaries from surrounding and interconnected tissues (i.e. fat body, muscle sheath, and oviduct) without damaging the ovarian cells. Thus, in order to minimize damage or death to ovarian cell types of interest, we elected to include these surrounding cell types in our analysis.

Female flies were selected on the day of eclosion and maintained at 25°C with access to males and yeast supplement for 3 days (a common experimental condition in many studies). Flies were then dissected in complete medium (Grace’s Insect Basal Medium supplemented with 15% FBS). To prevent cell clumping, ovaries were transferred to a tube containing 300 *µ*l EBSS (no calcium, magnesium, and phenol red) and gently washed for 2 minutes. The EBSS was then removed and the tissue was dissociated in 100 *µ*l Papain (50 U/mL in EBSS and previously heat activated in 37°C for 15 minutes) for 30 minutes. The suspension was mechanically dissociated every 3 minutes by gentle pipetting up and down. To quench the digestion, 500 *µ*l complete medium was added to dissociated cells. The suspension was then passed through a 40 *µ*l sterile cell strainer and centrifuged for 10 minutes at 700 RCF to remove large, undissociated eggs (with eggshell) and debris. This also filtered out larger germline cells which increase dramatically in size around stage 9 [53]. Supernatant was removed and single cells were re-suspended in 100 *µ*l. Cell viability was assayed using Trypan Blue and estimates of cell concentration were made using a hemocytometer. Cells were then further diluted to an approximate, final concentration of 2,000 cells/*µ*l according to 10X Genomics recommendations.

#### 10X Genomics library preparation

Single-cell libraries were prepared using the Single Cell 3’ Library & Gel Bead Kit v2 and Chip Kit according to the recommended 10X Genomics protocol. Single cell suspension was loaded onto the Chromium Controller (10X Genomics). Library quantification assays and quality check analysis was performed using the 2100 Bioanalyzer instrument (Agilent Technologies). The library samples were then diluted to a 10nM concentration and loaded onto two lanes of the NovaSeq 6000 (Illumina) instrument flow cell for a 100-cycle sequencing run. A total of 429,855,892 reads were obtained for the sample, with 28,995 mean reads per cell.

### Quantification and statistical analysis

#### Pre-processing Chromium single-cell RNA-seq output

The raw sequencing data for the 10X Genomics Chromium single-cell 3’ RNA-seq library were initially processed using Cell Ranger (version 3.0.0), the recommended analysis pipeline from the Chromium single-cell gene expression software suite. The reference index for Cell Ranger was built using the *Drosophila melanogaster* Release 6 reference genome assembly [80] made available on the Ensembl genome database. The cellranger count pipeline for alignment, filtering, barcode counting and UMI counting was used to generate the multidimensional feature-barcode matrix of 14,825 cells (S1 Fig).

The Cell Ranger output was used for further processing using the R package Seurat (v2.3.4) [12, 84]. As part of this processing, multiplet cells (those with less than 775 genes expressed per cell; setting a maximum of 2200 genes, and 18,000 UMIs per cell) and dead cells (greater than 1% mitochondrial gene expression) were filtered from the dataset (S1 Fig). Feature counts were log-normalized and scaled using default options (S1 Fig). Unwanted sources of intercellular variability were removed by regressing possible variation driven by number of UMIs and mitochondrial gene expression during data scaling. Scores for the expression of an expansive list of *Drosophila* G2/M and S phase genes (S2 File) were assigned to each cell which enabled the calculation of the difference between G2/M and S phase scores, using the function CellCycleScoring. This cell cycle score was then regressed from the downstream analysis to maintain the signals separating dividing and non-dividing cells but eliminating subtle differences among proliferative cells. Based on this score, the cells were assigned a cell cycle phase (S2 Fig). To assemble these cells into transcriptomic clusters using meaningful features, the number of random variables in our dataset was reduced by obtaining sets of principal component (PC) vectors. Significant PCs were obtained by performing Principal Component Analysis (PCA), using 897 highly variable genes as input. The first 30 significant PCs were selected based on the Elbow method as input for UMAP clustering using default parameters. Altogether, these pre-processing steps resulted in a primary UMAP of 12,671 cells (S1 Fig).

#### Manual removal of contaminated cells using biological markers

The clusters obtained in this primary UMAP construction were further processed for ambient RNA contamination removal (cleaned) based on aberrant gene expression patterns. Since we did not find any unique cluster for ovary/oviduct associating neuronal cell types (expressing commonly known neuronal cell markers *elav*), all cells expressing *elav* were considered contaminant and removed from the dataset. Similarly, we cleaned the germline clusters by removing cells that expressed somatic cell markers: *dec-1*, *Yp1/2/3*, *psd*, *Vml*, *Vm32E*, *Vm26Ab*, and *tj*; adipocyte marker: *Ilp6*; muscle cell markers: *Zasp66* and *Mp20*; and hemocyte marker: *Hml*. We cleaned the early somatic, polar, stalk, and mitotic follicle cell clusters by removing cells expressing germline cell markers: *osk* and *bru1*; mid-late somatic cell markers: *dec-1*, *Vm32E*, *Vm26Ab*, and *psd*; and hemocyte marker: *Hml*. We cleaned the mid-late clusters for cells expressing germline markers: *osk* and *bru-1*; muscle cell markers: *Zasp66* and *Mp20*; adipocyte marker: *Ilp6*; and hemocyte marker: *Hml*. We cleaned the muscle cell clusters by removing cells expressing germline cell markers: *osk* and *yl*; somatic cell markers: *tj*, *Yp1/2/* 3, *Vm32E*, *Vm26Ab*, *dec-1*, *psd*, and *Vml*; and hemocyte marker: *Hml*. We cleaned the adipocyte cluster by removing cells expressing somatic cell markers: *tj*, *dec-1*, *psd*, *Vml*, *Vm32E*, *Yp1/2/3*, and *Vm26Ab*; germline cell markers: *osk* and *yl*; muscle cell markers: *Zasp66* and *Mp20*; and hemocyte marker: *Hml*. A cut-off value of greater than 2 logFC was used to remove the contaminant cells. This manual cleaning strategy resulted in an increased resolution in the total number of highly variable genes (limits: *>*0.4 dispersion; *>*0.01 and *<*3 average expression) from 897 to 1075 which were then used as input for PCA on the cleaned dataset. The final dataset of high quality cells consisted of 7,053 cells and 11,782 genes (S1 Fig).

#### Cluster Validation of Replicate Data by Canonical Correlation Analysis (CCA)

The final 7,053-cells dataset was further compared to a 1,521-cell biological replicate dataset to assess the fidelity of the clustering (especially the trajectory of follicle cell clusters). This replicate dataset was derived from an original unprocessed dataset of 2,148 cells with 11,791 genes that was passed through a less stringent filtering criteria (due to the low number of cells) of 250 genes per cells as a lower threshold, and a higher threshold of 900 genes per cell, 2000 UMIs and 1% mitochondrial gene expression. The two datasets were aligned using 2,926 genes with the highest dispersion in both datasets. To detect common sources of variation between the two datasets, Canonical Correlation Analysis (CCA) was performed and 75 correlation vectors were used for downstream clustering. Upon plotting the UMAP using both the datasets, we were able to validate the perceived trajectory of the follicle cells. All follicular-cell states and clusters obtained in the 7,053-cells dataset were recapitulated in the UMAP using both replicates. We only used the replicate datasets to validate the clustering analysis and did not use this dataset for further downstream analysis since the cell sampling varied and we were unable to achieve a comparable sequencing depth (median genes per cell for the 1,521-cells dataset is 404) between the two datasets. The larger dataset (replicate 2) was used for all downstream analysis (S1 Fig).

#### UMAP clustering analysis

The 7,053 cells dataset (replicate 2) was log-normalized and scaled again using default parameters. The 1075 highly variable genes were selected as input for PCA and the first 75 PCs were selected to build the Shared Nearest-Neighbor (SNN) graph for clustering. To assemble cells into transcriptomic clusters, graph-based clustering method using the SLM algorithm [8] was performed in Seurat. We chose to plot clusters on a UMAP (Uniform Manifold Approximation and Projection) because this dimensionality reduction technique arranges cells in a developmental time-course in a meaningful continuum of clusters along a trajectory [6]. A number of resolution parameters, ranging from 0.5 to 6 were tested which resulted in 14 to 46 clusters. The relationship between clusters in each resolution was assessed using the R package clustree [103], based off of which a resolution of 6 was selected to obtain an initial number of 46 clusters (S2 Fig). Differentially expressed markers specific to each cluster were identified using the function FindAllMarkers (S3 File) and clusters with no unique markers were merged with their nearest neighbor after careful consideration of the differences in average expression pattern in each cluster. The final number of clusters was decided based on the uniqueness of observed and expected gene markers and the relative relationships with other clusters (S2 Fig). Cell type identities were then assigned to each cluster using known (S1 File) and experimentally validated markers.

#### Unsupervised re-clustering of cell subsets using Monocle (v2)

Smaller subsets of cells from the entire dataset were selected using the SubsetData function in Seurat. These subsets were re-clustered and imported into Monocle (v2) [74, 93] for further downstream analysis using the importCDS() function, with the parameter import_all set to TRUE to retain cell-type identity in Seurat for each cell. The raw UMI counts for these subsetted datasets were assumed to be distributed according to a negative binomial distribution and were normalized as recommended by the Monocle (v2) pipeline. The number of dimensions used to perform dimensionality reduction was chosen using the Elbow method. The cells were clustered in an unsupervised manner using the density peak algorithm where the number of clusters was set for an expected number of cell types (as in for early follicle cell differentiation states) or cell states (as in mitotic-endocycle transition state, along with mitotic and endocycling follicle cells). The number of cell clusters, in case of the “germline cells” subset and the “oviduct cells” and “muscle cells” subset was chosen in an unsupervised manner based on significant rho (local density) and delta (distance of current cell to another cell of higher density) threshold values.

#### Pseudotime Inference Analysis and Identification of Lineage-Specific Genes of Interest

Pseudotime inference analysis on known cell differentiation programs of oogenesis was performed using Monocle (v2). Cells were ordered in an unsupervised manner on a pseudotemporal vector based on genes that are differentially expressed over pseudotime between cell type identities assigned in Seurat or cell states identified as clusters in Monocle, depending on the clustering as mentioned in the previous section. Lowly expressed aberrant genes were removed from the ordering genes. Multiple trajectories were generated by ordering the cells using different numbers of statistically significant (q*<*0.05) genes that are expressed in a minimum number of predetermined cells, and the efficacy of the trajectories was tested with validated marker gene expression. The trajectory that reflected the most accurate cell state changes was then selected for downstream analysis. To assess transcriptional changes across a branching event, as seen in the early somatic and the polar/stalk trajectories, the function BEAM was used to analyze binary decisions of cell differentiation processes across a branch.

#### Gene ontology (GO) term enrichment Analysis

Genes were selected for downstream GO term enrichment analysis from the pseudotemporal heatmap by cutting the dendrogram that hierarchically clustered the genes expressed in a similar pattern across pseudotime using the R based function cutree [7]. The web-based server g:Profiler [75] and PANTHER [65] were then used for functional enrichment analysis on the genes. A user threshold of p=0.05 was used for these analyses.

## Results

### ScRNA-seq identifies unique cell clusters and markers to assign cell type identities

We generated the scRNA-seq library from a cell suspension of freshly dissected ovaries (and connected tissues) from adult female flies (Fig 1A). Following library sequencing, extensive quality control, and cell type-specific marker validation, we recovered 7,053 high-quality cells and clustered them into 32 cell-type identities (Fig 1B, S1 Fig and S2 Fig). This dataset has an average of ∼7,100 UMIs and ∼1,300 genes per cell, with each cell type having variable levels of mRNA content and gene expression (Fig 1C and 1D). We plotted this dataset on a scale of two primary axes for visualization using Uniform Manifold Approximation and Projection (UMAP) for dimension reduction of the cell/gene expression matrix (Fig 1B). This UMAP reflects the temporal and spatial development over the entirety of oogenesis, with connected ovarian clusters forming linear trajectories from stem cells onward, while surrounding tissues with non-temporally transitioning cells (muscle sheath, oviduct, adipocytes, and hemocytes) arranged in compact and isolated clusters (Fig 1B and S2 Fig).

**Fig 1.**
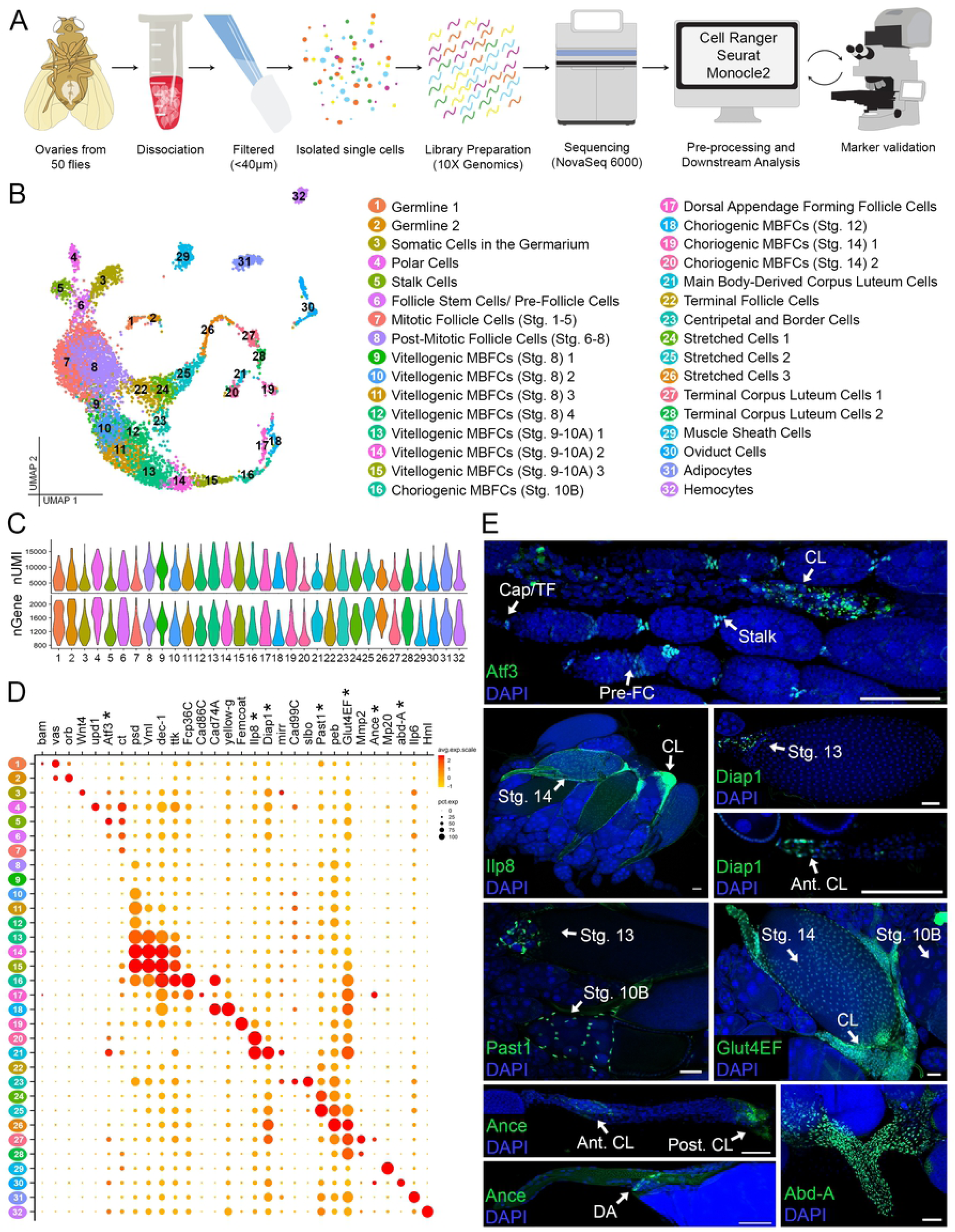
ScRNA-seq of adult *Drosophila* ovary and interconnecting tissues. (A) Illustration of the overall workflow (See also S1 Fig). (B) Annotated UMAP of 7,053 high-quality cells grouped into 32 semi-supervised clusters and labelled according to cell type and stage (See also S2 Fig and S2 File). MBFCs=Main Body Follicle Cells. (C) Number of UMI (nUMI) and genes (nGene) per cluster. (D) Dot plot of identifying marker genes (See also S1 File). Newly identified marker genes are indicated (*). (E) Experimental validation of the 7 new marker genes shown in D. All expression (green) is marked using GFP-tagged proteins under endogenous control except Ance, marked using RFP under T2A-Gal4 control. All images are z-projections. Additional cell-type and stage-information is indicated (Cap/TF= Cap and Terminal Filament Cells, Pre-FC= Pre-Follicle Cells, Stalk= Stalk Cells, CL= Corpus Luteum Cells, Stg.=Stage, Ant. CL= Anterior Corpus Luteum Cells, Post. CL= Posterior Corpus Luteum Cells, DA= Dorsal Appendage Forming Follicle Cells). DAPI marks nuclei. Scale bar = 50 *µ*m.

Established cell-type and stage-specific markers were used to identify the majority of the clusters (S1 File and Fig 1D). For the remaining clusters with no known markers, we assigned identity using expression patterns of at least 7 newly validated genes (Fig 1D and 1E). *Atf3* and *abd-A* were used to identify cell types such as stalk cells and oviduct cells. *Past1* was used to identify the stretched cells, and *Ilp8*, *Diap1*, *Glut4EF*, and *Ance* were used to identify late-staged follicle cells. Most of the new markers have overlapping expression in multiple cell types. For example, *Atf3*, a transcription factor involved in lipid storage [78], marks the cap and terminal filament cells in the germarium, pre-follicle cells, stalk cells, and corpus luteum cells (Fig 1E). Similarly, some markers are expressed in cells across multiple timepoints, thus marking a single cell type in several clusters. For example, *Past1*, which encodes a plasma membrane protein known to interact with Notch, marks the stretched cell lineage in clusters 24, 25, and 26 [72]. Altogether, we were able to assign cell type identities for all clusters and identified 6,296 genes that show significant expression in different clusters. Among them, 828 are unique markers for clusters, that may be potentially specific to individual cell-types (S3 File).

### The transcriptional patterns of early germline development

Oogenesis begins in the germarium at the most anterior tip of each ovariole. There, supported by somatic niche cells, two to three germline stem cells (GSCs) produce daughter cells which move posteriorly through the niche and differentiate into cystoblast cells (CCs) [16]. These cells undergo four more rounds of synchronized mitosis with incomplete cytokinesis, producing 16 interconnected germline cyst cells. One of these cells becomes a transcriptionally quiescent oocyte, while the others develop into nurse cells that synthesize and transport products into the oocyte through ring canals [22](Fig 2A).

**Fig 2.**
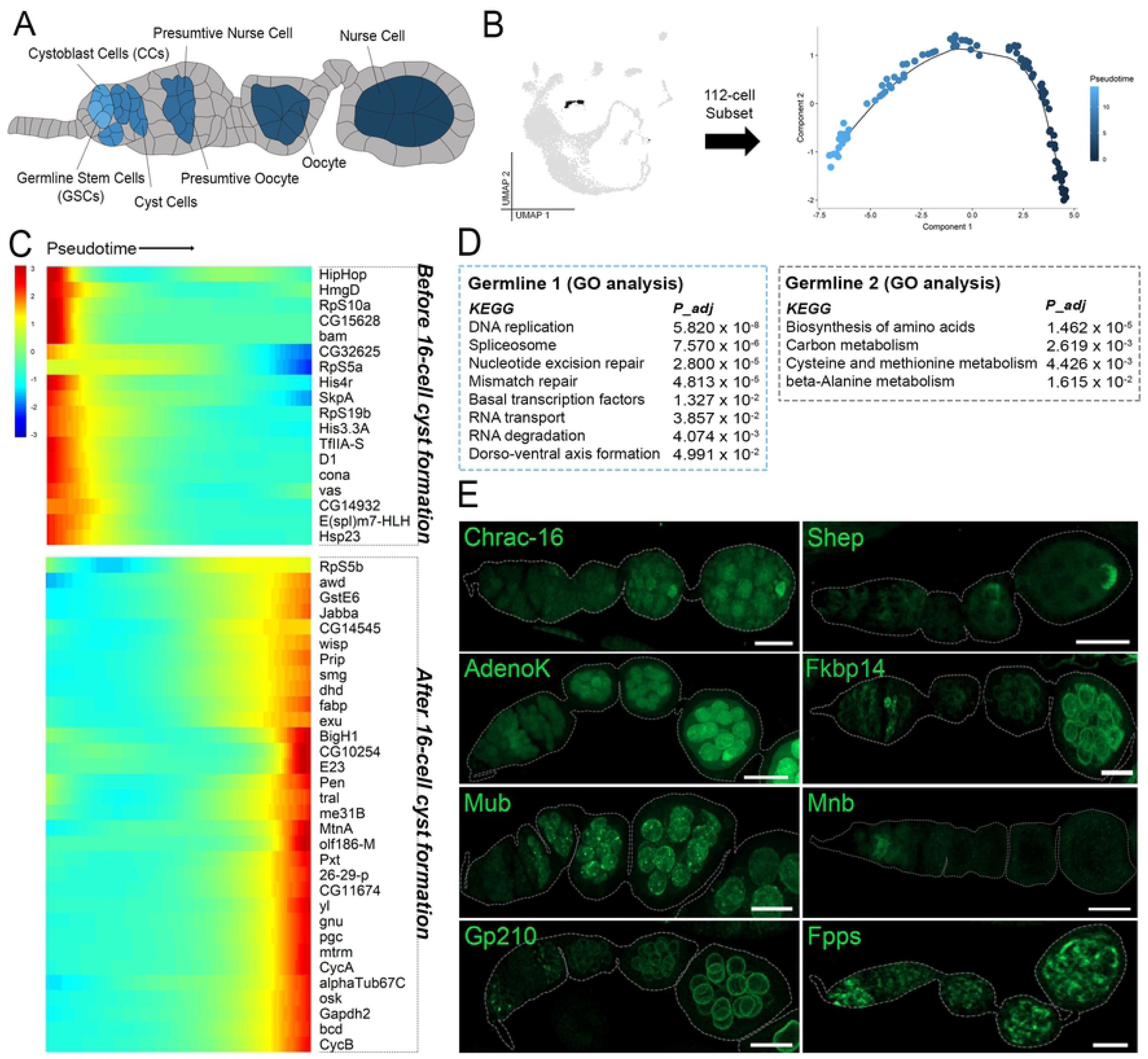
Expression patterns of germline cells during early development. (A) Illustration of early oogenesis featuring annotated germline cell types of interest (colored according to pseudotime inference in B) and somatic cells (grey). (B) Fig 1B UMAP (grey) at left highlighting the 112-cell subset of germline clusters 1-2 (black) re-clustered in Monocle for pseudotime analysis. Subset tSNE plot at right with pseudotime scale. (C) Pseudotime-ordered heatmap of expression from before and after 16-cell cyst formation. Minimum expression = 5 cells; *q <* 1*e*^−5^. (D) KEGG pathway terms and enriched for germline 1 (blue box) and germline 2 (black box) clusters. Adjusted p-values (*P_adj_*) are provided for each term. (G) Validation for germline expression (green) using GFP-tagged proteins under endogenous control. All images are z-projections. Ovarioles are outlined in grey. Scale bar = 20 *µ*m.

The germline cells in our dataset were size selected through manual filtration (see Materials and methods), resulting in a sampling from GSCs to those in mid-oogenesis. These cells form a two-cluster trajectory (Fig 1B). The Germline 1 cluster includes cells in region 1 of the germarium (marked by *bam* expression) and the Germline 2 cluster includes cells from region 2 of the germarium and onward (marked by *orb* expression) [55, 63] (Fig 1D). The formation of the 16-cell cyst occurs at the boundary of germarium region 1 and 2. To uncover the underlying expression changes occurring at this time, we arranged the 112 germline cells on a pseudotemporal axis (Fig 2B) and plotted the differentially expressed genes along pseudotime. This revealed 50 genes that are expressed significantly before or after 16-cell cyst formation (Fig 2C). Gene Ontology (GO) enrichment of KEGG-pathway terms across pseudotime revealed the broad differences in activity before and after 16-cell cyst formation. Germline 1 cells are enriched for DNA replication and repair genes and Germline 2 cells switch to an enrichment in biosynthetic- and metabolic-pathway genes (Fig 2D). This is strikingly similar to the recent findings in a testis scRNA-seq study, which suggest an increase in mutational load in the immature germline cells of the testis and an early expression bias for DNA repair genes [97].

Selected germline-specific genes were experimentally validated and show varying expression patterns in the early stages of oogenesis (Fig 2E). Among these newly identified germline markers, specific expression of *Mnb*, a Ser/Thr protein kinase, in region 1 of the germarium and *Mub*, an mRNA splicing protein which appears only after 16-cell cyst formation, is of special interest [73, 89]. Other interesting expression patterns were identified in genes such as *Fpps* and *Gp210*, which briefly appear in the germarium, disappear for several for several for several aring, demonstraesng the dynamic regulation of early germline cell transcription.

### Transcriptional trajectory of early somatic differentiation

The anterior region of the germarium houses somatic cells that include eight to ten terminal filament cells, a pair of cap cells, and the escort, or inner germarium sheath (IGS), cells. These collectively form the germline stem cell niche [31, 101] (Fig 3A). In the next region of the germarium is the somatic stem cell niche where two or more follicle stem cells (FSCs) differentiate to form the pre-follicle cells (pre-FCs) that envelope the germline cyst cells. As egg chambers pinch off from the germarium, pre-follicle cells at the two poles assume polar cell fate upon Notch activation. The anterior polar cells then promote the specification of the stalk cells through JAK/STAT signaling [3]. The polar and stalk cells cease division upon differentiation while the other follicle cells remain mitotically active [82].

**Fig 3.**
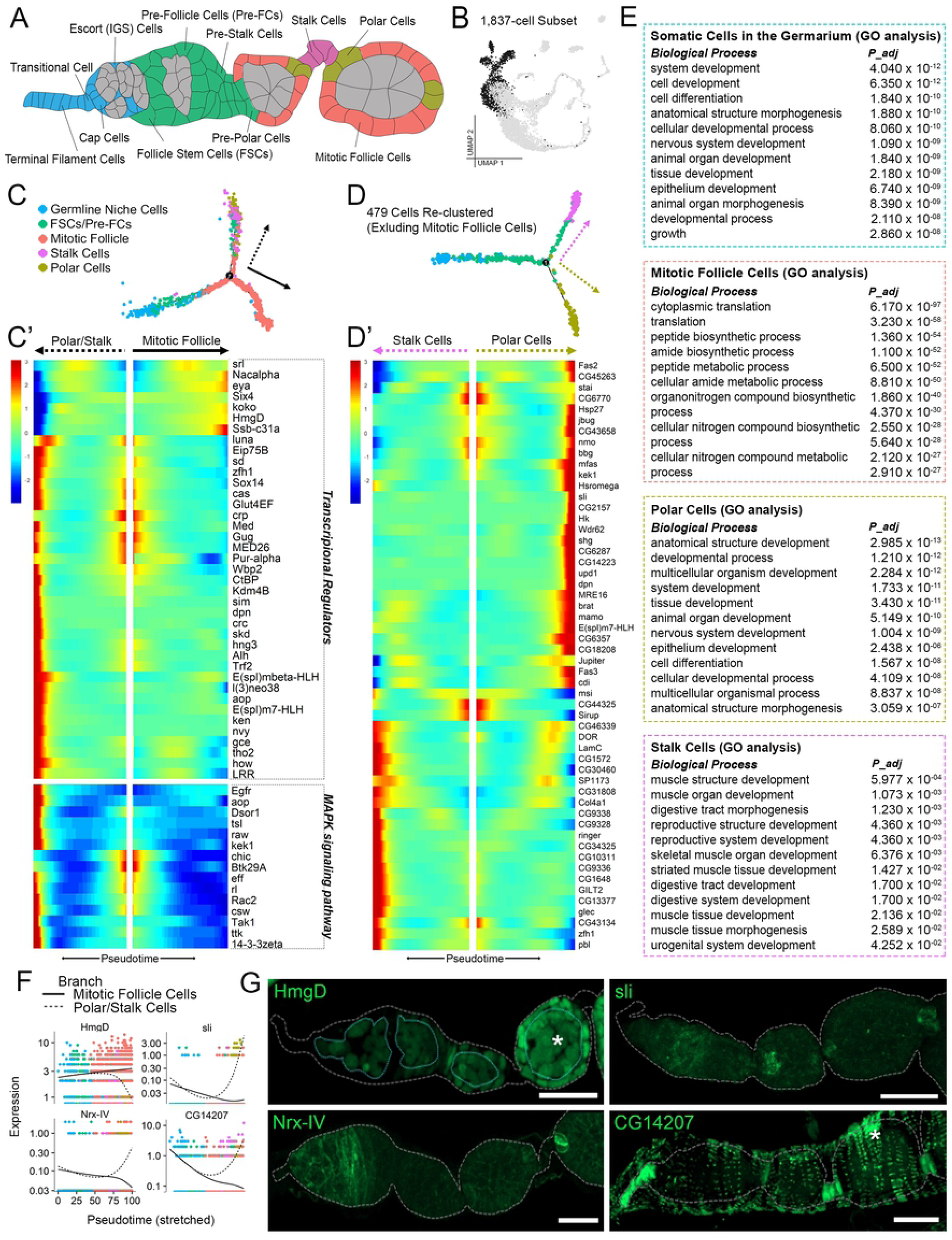
Transcription of early somatic cells during differentiation. (A) Illustration of early oogenesis featuring annotated somatic cell types of interest (colored according to identity in C) and germline (grey). (B) Fig 1B UMAP (grey) highlighting the 1,837-cell subset of early somatic cell clusters 3-7 (black) re-clustered in Monocle for pseudotime analysis. (C) Trajectory tSNE of subset cells ordered along pseudotime (C’) Pseudotime-ordered heatmap from trajectory in C with select genes (transcriptional regulators: GO:0140110 or PC00218, and MAPK signaling pathway: KEGG:04013) selected from expression in a minimum of 20 cells, q*<*0.05. (D) Trajectory tSNE of the 479-cell subset (excluding mitotic follicle cells). (D’) Pseudotime-ordered heatmap from trajectory in D. Minimum expression = 20 cells, *q <* 1*e*^−5^. (E) Enriched Biological Process terms for somatic cells in germarium cluster and mitotic follicle, polar, and stalk cell branches. Adjusted p-values (*P_adj_*) are provided for each term. (F) Expression plots of validated genes arranged along pseudotime (from trajectory in C) comparing the mitotic follicle cell (solid line) and polar/stalk cell (dotted line) branches. (G) Experimental validation of select genes (green) using GFP-tagged proteins under endogenous control. All images are z-projections. Ovarioles are outlined in grey. Germline outlined in top left image. Some expression is also observed in other cell types and marked with an asterisk (epithelial sheath cells in bottom right image and germline cells in top left image). Scale bar = 20 *µ*m.

Due to the unsupervised nature of our clustering, the somatic cells in the germarium are cluster together Fig 1B). This suggests a common transcriptomic signature which may be a response to the shared stem cell niche signaling. GO analysis for this group revealed an unexpected enrichment of nervous system development related genes, among more general development- and morphogenesis-related genes (Fig 3E).

To determine the transcriptional trajectory during early somatic differentiation, we arranged the 1,837-cell subset from clusters containing somatic cells of the germarium, polar cells, stalk cells, and mitotic follicle cells on a pseudotemporal axis (Fig 3B and 3C). This pseudotemporal trajectory establishes a divergence of the follicle cell lineage after FSC/pre-FC differentiation, as the branch for mitotic follicle cells separates out from a common branch for the polar/stalk cell lineage (Fig 3C). This trajectory is consistent with the notion that polar and stalk cells share a common precursor stage and share expression of certain commonly upregulated transcription factors as shown in other studies [18, 95].

Considering the importance of transcriptional regulation in differentiation, we analyzed the temporal patterns of highly expressed genes selected for their function as either transcription regulators (GO:0140110) or transcription factors (PC00218) (Fig 3C’). Plotting these genes across pseudotime revealed that the polar/stalk cell fates are transcriptionally dynamic, involving genes from many signaling pathways. We highlighted the genes involved in the MAPK pathway (Fig 3C’). Fewer transcription factors are expressed in the mitotic follicle cell lineage (Fig 3C’). Among them are the chromatin remodeling protein HmgD and its physical interactor Nac*α*, suggesting a role of epigenetic regulation in the proliferative effort of these cells [34, 39] (Fig 3C’, 3F and 3G). The mitotic follicle cell lineage also shows a differential enrichment of ribosomal genes (*KEGG* : 03010*, P_adj_* = 2.20*e*^−49^), probably to support the upregulation of biosynthetic processes to sustain rapid proliferation (Fig 3E).

### Fate decisions during polar and stalk cell differentiation

To characterize the fate separation between polar and stalk cells, we excluded the mitotic follicle cells from further analysis. The resulting 479 cells were then ordered once again along a pseudotemporal axis (Fig 3D). The resulting trajectory shows that the polar cells differentiate earlier than the stalk cells, which is consistent with the evidence that chemical cues from polar cells initiate stalk cell differentiation [3, 95]. To further identify genes that regulate polar and stalk cell differentiation, we plotted the most significant (*q <* 1*e*^−5^) differentially-expressed genes between the two fates (Fig 3D’). GO analysis of biological functions in the polar cell branch revealed a remarkable number of genes involved in processes related to nervous system development, neurogenesis, and neuron differentiation, similar to neuron-related expression in somatic cells of the germarium (Fig 3E).

Many such genes (e.g., *Fas2*, *bbg*, *kek1*, *sli*, *shg*, *brat*, *Fas3*, and *CG18208*) produce junctional proteins (*CG* : 0005911*, P_adj_* = 5.563*e*^−4^) or proteins at the cell periphery (*CG* : 007194*, P_adj_* = 2.568*e*^−2^) (Fig 3D’) We validated the expression of *sli*, a novel polar cell marker, which is a secreted ligand for the Slit/Robo signaling pathway (Fig 3F and 3G). Another validated polar cell marker, *Nrx-IV*, is also associated with this pathway [5](Fig 3F and 3G). In addition to axon guidance in developing neurons, Slit/Robo has been implicated in the regulation of tissue barriers [98], which is consistent with the observation that polar cells are terminally differentiated barriers between each egg chamber unit and connecting stalk cells [37].

GO term analysis of stalk cell specific genes indicates a highly significant (*q <* 1*e*^−5^) upregulation of extracellular matrix genes (e.g. *Col4a1*, *LanB1*, and *vkg*) and cytoskeletal genes (e.g. *LamC* and *βTub56D*) that are also involved in muscle structure development (Fig 3D’ and 3E). Supporting this finding, we found a novel stalk cell marker *CG14207*, that is also expressed in epithelial muscle sheath (Fig 3F and 3G). Its human homolog, HspB8, interacts with Stv at the muscle sarcomere as part of a chaperone complex required for muscle Z-disc maintenance [2].

### Catalytic genes upregulated during Mitosis-Endocycle transition of follicle cells

The transition between early and middle oogenesis (stages 6-7), occurs when the germline cells upregulate the ligand Dl, activating Notch signaling in the follicle cells, which initiates a mitosis-endocycle (M/E) switch [26] (Fig 4A).

**Fig 4.**
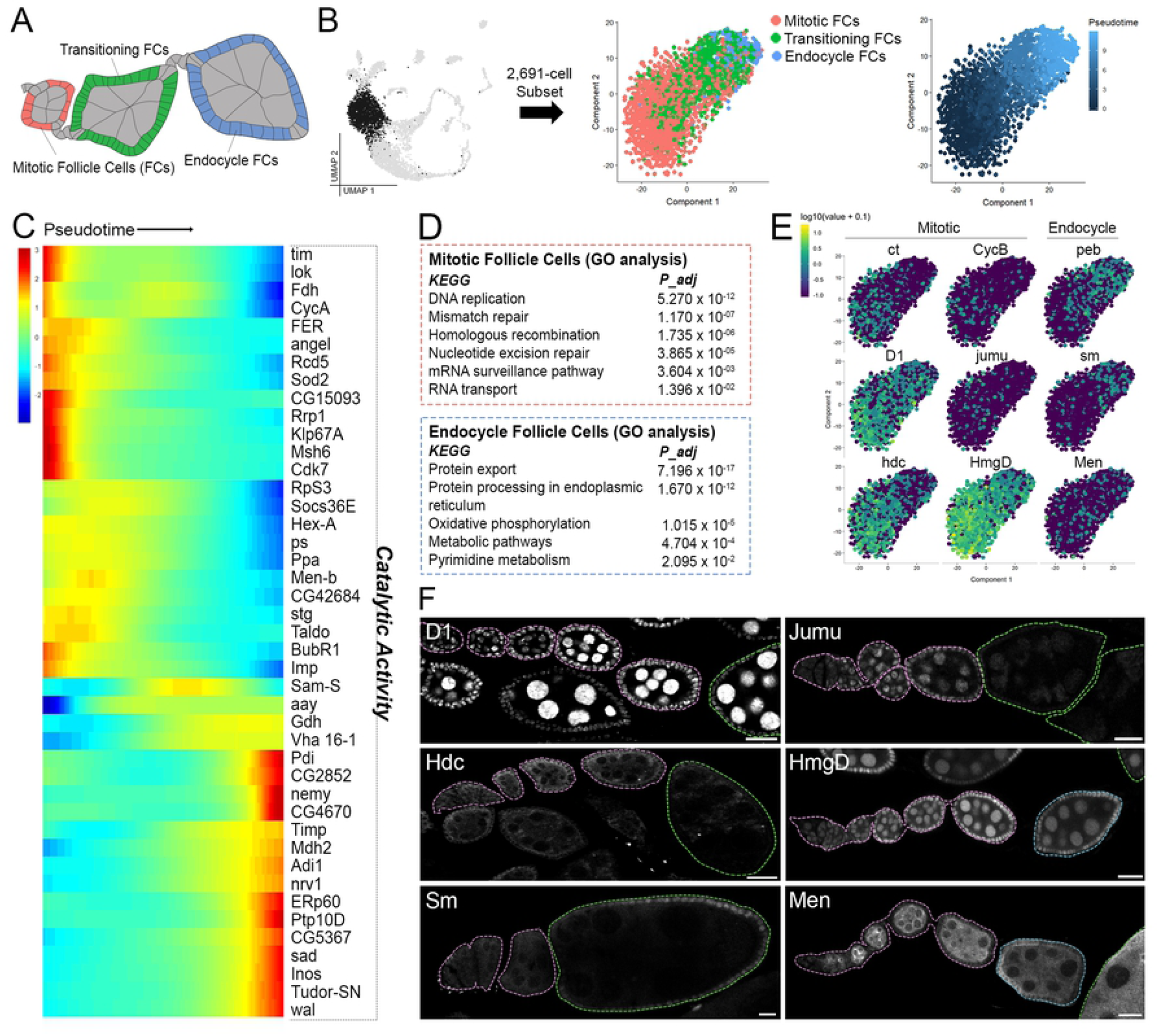
Gene expression during Mitotic-Endocycle transition in follicle cells. (A) Illustration of follicle cells of interest during M/E switch (colored according to their cluster color in B) with all other cells in grey. (B) Fig 1B UMAP (grey) highlighting the 2,691-cell subset of early to mid-staged follicle cells from clusters 7-8 (black) re-clustered in Monocle for pseudotime analysis (left). Subset tSNE with cluster annotation informed by ct, CycB, and peb marker expression shown in E (center). Subset tSNE with pseudotime colors (right). (C) Pseudotime ordered heatmap of highly expressed genes grouped by catalytic activity (GO:0003824). Minimum expression = 20 cells; q=0.05. (D) KEGG pathway terms enriched in mitotic and endocycling follicle cells (early and late expressing genes respectively from C). (E) Feature plots for select genes showing differential patterning in either mitotic or endocycle follicle cells. Top row genes (*ct*, *CycB*, and *peb*) are known markers. The others are newly identified. (F) Experimental validations for newly identified M/E switch markers (white) using GFP-tagged proteins under endogenous control. Ovarioles are outlined and colored according to stage: germarium and mitotic stages (pink), transitional stage (green), and endocycle stages (blue). All images are a z-slice through the center of each ovariole. Scale bar = 20 *µ*m.

To understand the regulation of the M/E switch at the single-cell level, we re-clustered the 2,691 follicle cells from clusters 7, 8, and 9 and arranged them across pseudotime (Fig 4B). Known Notch targets were used to validate cluster identity: *ct* and *CycB* in mitotic cells, *peb* in endocycling cells [85, 86], and all three in transitioning cells (Fig 4E). Pseudotime analysis revealed a linear arrangement for genes that change expression levels during the M/E switch. We validated some of these newly identified genes. For example, *D1*, *jumu*, and *hdc*, are down-regulated, while *Men* and *sm*, are upregulated in post-mitotic follicle cells (Fig 4F). The NADP[+] reducing enzyme, *Men*, is upregulated significantly in the anterior follicle cells and has a membrane localization. *Sm*, a member of the heterogeneous ribonucleoprotein complex is of special interest given its ability to regulate Notch activity during wing development [50]. Its enrichment in endocycling follicle cells suggests a potential role for sm in Notch-mediated M/E switch. Noticeably, upon GO term enrichment analysis of all significantly expressed genes that change as a function of pseudotime during the M/E switch, we found 43 genes with catalytic activity (GO:0003824) (Fig 4C). Enriched KEGG-pathway-related terms reveal an expression bias for proliferation and DNA repair associated genes in mitotic follicle cells, whereas endocycling cells express protein-processing and metabolic genes (Fig 4D).

### Transcriptomic divergence of mid-staged follicle cells with subsequent convergence

During early oogenesis, access to morphogen signals from polar cells are restricted to the nearby terminal follicle cells (TFCs) on either end of the egg chamber [42]. The posterior terminal follicle cells receive a signal from the oocyte to activate EGFR signaling around stage 6, marking a symmetry breaking event in follicle cells. Cells at the anterior terminal further specify into border, stretched, and centripetal cells and undergo massive morphological changes during stages 9-10B [100] (Fig 5A).

**Fig 5.**
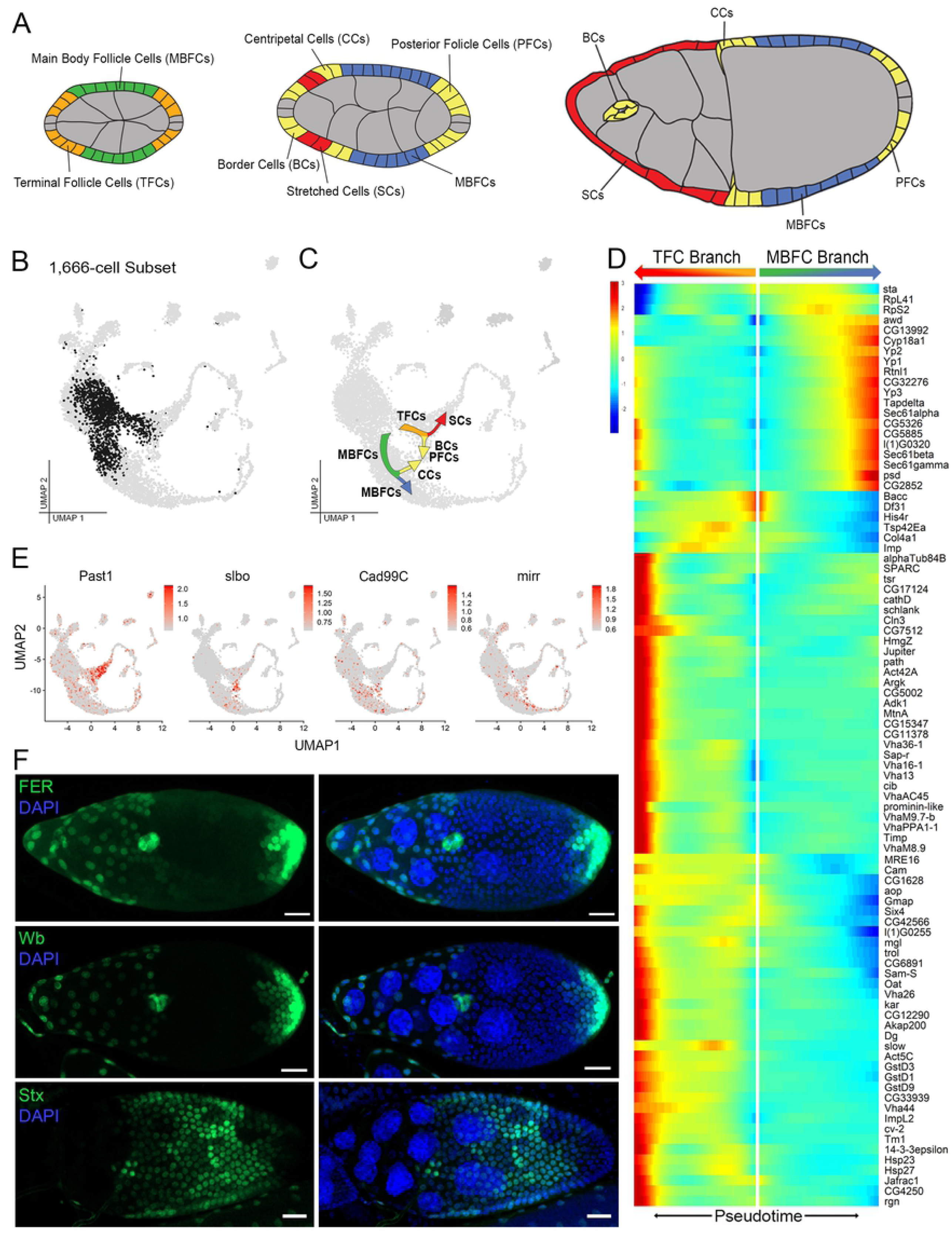
Transcriptional divergence of terminal follicle cells during symmetry breaking with subsequent convergence of slbo-expressing cells. (A) Illustration of annotated follicle cell types during symmetry breaking and differentiation (colored by type) with all other cell types shown in grey. Stalk cells not shown. (B). Fig 1B UMAP (grey) highlighting the 1,666-cell subset of mid-staged follicle cells in clusters 8-10 and 22 (black) re-clustered in Monocle for pseudotime analysis. (C) Fig 1B UMAP (grey) annotated with cell type lineage information based on markers in E. (D). Pseudotime ordered heatmap of gene expression during the Terminal Follicle Cell (TFC) and Main Body Follicle Cell (MBFC) branching in C. Minimum expression = 20 cells; *q <* 1*e*^−20^. (E). Feature plots of marker genes used for identification in C. Past1= stretched cells (SCs), slbo= border cells (BCs), polar follicle cells (PFCs), and centripetal cells (CCs), Cad99C= centripetal cells (CCs), mirr= main body follicle cells (MBFCs). (F) Experimental validation of select gene expression (green) in cells after symmetry breaking (not shown in heatmap in D). All lines express GFP under T2A-Gal4 control for each gene. *FER* and *wb* are expressed in SCs, BCs, and PFCs. *Stx* is expressed in main body follicle cells. All images are z-projections. DAPI marks nuclei. Scale bar = 20 *µ*m.

Our dataset shows an unanticipated transcriptomic divergence for post-mitotic follicle cells, which provides a transcriptional basis for follicular symmetry breaking (Fig 1B). To identify the fate assumed by the cells in each resulting branch, we validated the expression of known markers at this stage and also novel markers uncovered from re-clustering 1,666 cells of this stage (Fig 5B). The main body follicle cell (MBFC) branch was identified using *mirr* and *Cad99C* expression [21, 49]. And the TFC branch identity was validated by the expression of newly-identified anterior terminal cell marker, *Past1* (Fig 5E).

We took the 1,666-cell subset of follicle cells during symmetry breaking and arranged them on a pseudotemporal axis (Fig 5B). Then we performed a GO term enrichment analysis of the differentially expressed genes at the branching point between MBFC and TFC fate. The MBFC fate shows an enrichment of genes in protein export (*KEGG* : 03060*, P_adj_* = 8.55*e*^−20^) and protein processing in the endoplasmic reticulum (*KEGG* : 04141*, P_adj_* = 1.13*e*^−17^); whereas the TFC fate has an enrichment of genes in endocytosis (*KEGG*04144*, P_adj_* = 1.70*e*^−9^), proteasome (*KEGG* : 03050*, P_adj_* = 3.46 7), phagosome (*KEGG*; 04145*, P_adj_* = 6.97*e*^−6^), glutathione metabolism (*KEGG* : 00480*, P_adj_* = 2.09*e*^−2^), oxidative phosphorylation (*KEGG*00190*, P_adj_* = 2.01*e*^−2^), and Hippo pathway (*KEGG* : 04391*P_adj_* = 3.95*e*^−2^). The 89 genes that show significant differences between these two branches along pseudotime are highlighted in a heatmap (Fig 5D). Many genes are differentially upregulated in these two branches much later in pseudotime.

We also identified novel genes showing expression that coincides with the symmetry breaking process (Fig 5F). These include *FER* and *wb*, which regulate cytoskeletal rearrangement, cell adhesion, and extracellular components. These genes may participate in cell shape changes necessary for border cell migration and/or stretched cell flattening [62, 68]. On the other hand, MBFC-specific expression of *stx* is interesting as it is involved with the proteasomal degradation regulating Polycomb (Pc) stability [29]. Maintenance of MBFC fate through regulation of chromatin modifiers is an attractive direction that merits further research.

### Expression profiles of migrating border and centripetal cells

During stages 9-10B, specialized subsets of TFCs transition from a stationary to migratory state. These include the border cells, which delaminate from the epithelium and move through the nurse cells to reach the oocyte. There, they meet the centripetal cells which migrate inward to cover the anterior end of the oocyte (Fig 6A).

**Fig 6.**
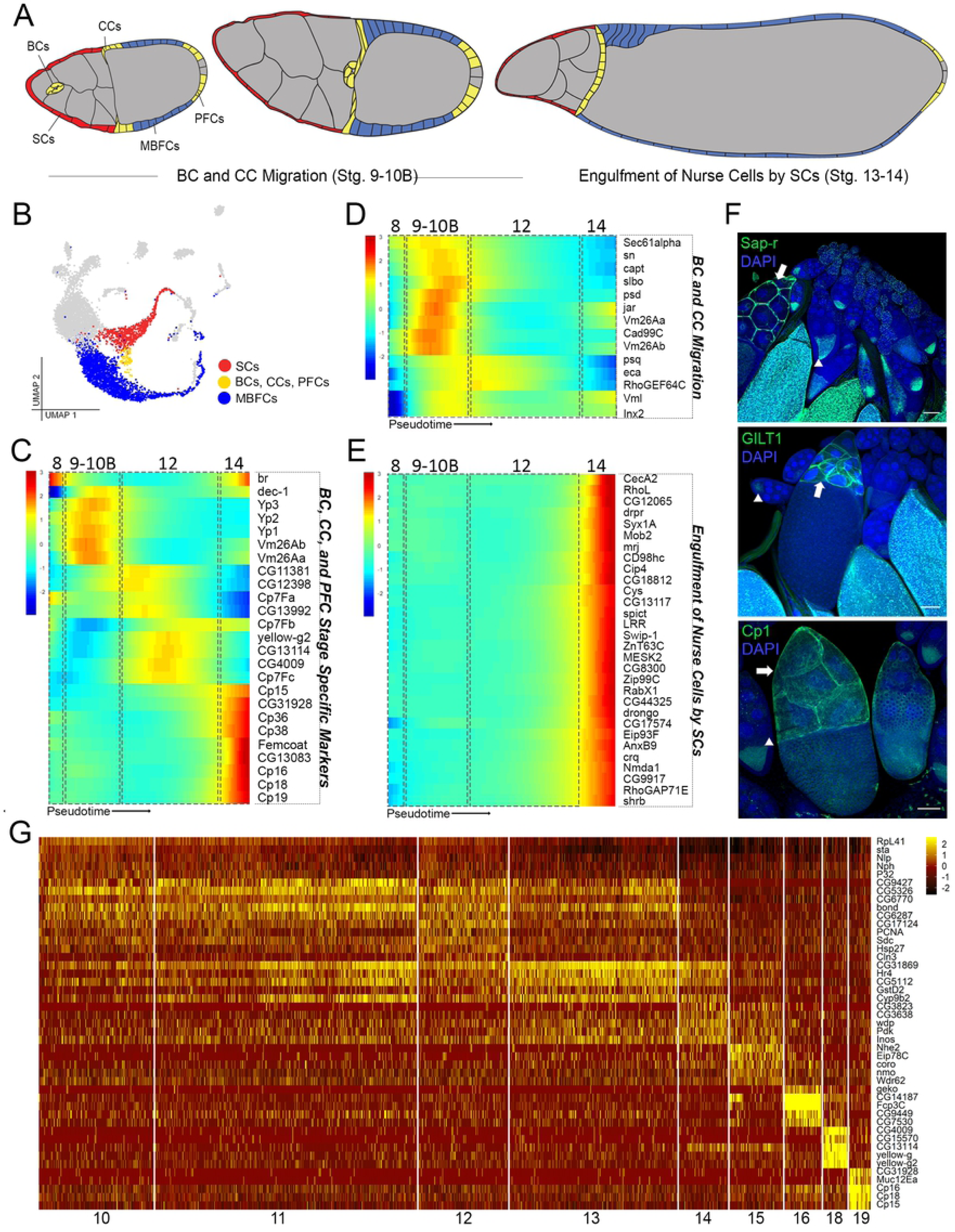
Gene expression in follicle cells during migration, nurse cell engulfment, and vitellogenesis. (A) Illustration of annotated follicle cells of interest (colored according to UMAP in B) with all other cell types in grey. Stalk cells not shown. (B) Fig 1B UMAP (grey) highlighting the mid-late stage follicle cell subsets re-clustered in Monocle for pseudotime analysis. Terminal follicle cell (TFCs) and stretched cell (SCs) subset = 798 cells from clusters 22-26 (red). Border cell, centripetal cell, and posterior follicle cell (BCs, CCs, PFCs) subset = 193 cells from cluster 23 (yellow). Main body follicle cell (MBFCs) subset = 1,988 cells from clusters 10-16 and 18-19 (blue). (C) Pseudotime ordered heatmap of stage 8-14 specific markers from red and yellow subsets from B. Estimated stage boundaries (dotted boxes) are superimposed on the heatmap. (D) Pseudotime ordered heatmap of genes during stage 9-10B (in cells from yellow and red subsets) with epithelial development genes (*GO* : 0060429*, P_adj_* = 1.101*e*^−5^) specifically highlighted. Minimum expression = 100 cells; q*<*0.05. (E) Pseudotime ordered heatmap of red and yellow subset genes in stage 14 highlighting the 30/79 genes also expressed in hemocyte cluster 32 from Fig 1B. Minimum expression = 50 cells; *q <* 0.05. (F) Experimental validation for three highly expressed genes in stretched cells (not shown in the heatmap in E) using GFP-tagged proteins under endogenous control. Arrows point to stretched cells and arrowheads point to additional expression in oocytes. All images are a single z-slice through the center of egg chambers. DAPI marks nuclei. Scale bar = 20 *µ*m. (G) Heatmap of top 5 highly expressed genes per cluster for the blue subset (clusters 10-16, 18-19 from Fig 1B).

In our plot, we found that the TFC and MBFC branches converge to form a distinct cluster marked by *slbo*, which is expressed in migrating border and centripetal cells [66] (Fig 5C). To examine the transcriptomic signature of these migratory cells, we first used known stage 8-14 markers [46, 92] to set stage boundaries for the TFC branch (Fig 6B and 6C). This boundary was then used to select gene expression specifically during cell migration. We highlighted 14 representative genes involved in epithelial development (*GO* : 0060429*, P_adj_* = 1.101*e*^−5^), the highly enriched GO term in this cluster. These include markers for border cell migration, such as *sn*, *jar*, and *Inx2* [25, 35, 45, 79]. We also detected in this cluster the expression of *Cad99C*, which has been reported in several main body follicle cells, and anterior-migrating centripetal cells [21]. These known markers confirm the correct selection of migrating cell types. This cluster also show expression of other stage 9-10B markers, such as vitelline membrane-related genes: *psd*, *Vm26Aa*, *Vm26Ab*, and *Vml* [30, 92, 106]. With the confidence in our selection of stage 9-10B migrating cells, we identified additional genes such as protein transmembrane transporter *Sec61α*, actin binding protein *capt*, cargo receptor *eca*, and Rho guanyl-nucleotide exchange factor *RhoGEF64C*, which may contribute to different aspects of the cell migration process [15, 34, 36, 41, 83] (Fig 6D).

### Stretched cells share the transcriptional signature with hemocytes as they engulf nurse cells

During the final stages of oogenesis (stages 13-14), after the nurse cells transfer their cytoplasm into the oocyte, the remaining nuclei and cellular contents are removed by the stretched cells. This phagocytic activity of stretched cells is reminiscent of the response of hemocytes upon infection [90]. To determine whether genes expressed in the stretched cell cluster are also expressed in hemocytes, we examined the stage 13-14 specific genes identified from the pseudotemporally arranged 798-cell subset of the TFC branch. We identified 11 genes in this cluster (*LRR*, *PGRP-SD*, *Irbp18*, *PGRP-LA*, *Hsp26*, *trio*, *bwa*, *Hsp67Bc*, *CecA2*, *Hsp27*, and *Hsp23*) categorized by their involvement in immune system process (GO:0002376). We also compared genes enriched in the stretched cells with those in the hemocyte cluster and found 79 genes in common. Of these, 30 genes with the highest expression are shown in a heatmap ordered across pseudotime (Fig 6E). Some immune genes have been identified previously in nurse cell engulfment, such as the phagocytic gene *drpr*, and a scavenger receptor gene *crq*, confirming sampling of the correct developmental time-point for analysis [64, 90]. The newly identified genes in the stretched cell cluster fall into six general categories of activity: endocytosis/vesicle mediated transport (*Syx1A*, *RabX1*, *AnxB9*, and *shrb*), antibacterial/immune response (*CecA1* and *LRR*), morphogenesis (*Mob2*, *CG44325*, *RhoGAP71E*, and *RhoL*), catalytic/metabolic (*CG12065*, *Cip4*, and *Nmda1*), lipid binding (*Cip4* and *Gdap2*), and metal ion transport, especially zinc and magnesium (*spict*, *Swip-1*, *ZnT63C*, and *Zip99C*). In addition, we validated three new stretched-cell genes (Fig 6F) which are also expressed in hemocytes: a proteolytic enzyme, *Cp1* involved in cellular catabolism, an oxidation-reduction enzyme, *GILT1*, involved in bacterial response, and *Sap-r*, a lysosomal lipid storage homeostasis gene with known expression in embryonic hemocytes [54, 81, 94]. Together, these findings suggest that stretched cells and hemocytes share transcriptomic signatures required for apoptotic cell clearance, reinforcing their role as “amateur” phagocytes at this stage of development [38].

### Gene expression of vitellogenic main body follicle Cells

The clusters for the MBFCs show an enrichment of genes that facilitate vitellogenesis (stages 8-14) and eggshell formation (stages 10-14; Fig 1D). We further analyzed the clusters of the MBFC clusters and found highly variable gene expression patterns (Fig 6B, and 6G). Genes enriched in clusters 10-13, presumably consisting of stage 8-10A MBFCs, include histone binding protein-coding genes such as *Nlp*, *Nph*, and *P32*, which have been shown to cooperate in the post-fertilization regulation of sperm chromatin [32]. Starting in cluster 16, marked by the stage 10B specific marker *Fcp3C*, chorion-related genes such as CG14187, acid phosphatase *CG9449*, and signaling receptor, *CG7530* show an upregulation. Stage-12 and 14 follicle cells (clusters 18 and 19 respectively) express well-known markers involved with chorion production (e.g. *CG4009*, *CG15570*, *CG13114*, *yellow-g*, *yellow-g2*, *CG31928*, *Muc12Ea*, *Cp16*, *Cp18*, and *Cp15*) [92] (Fig 6G).

### Cellular heterogeneity and markers in the corpus luteum

Ovulation occurs when a mature egg sheds the follicle-cell layer and exits the ovary on its way to be fertilized, following *Mmp2* -dependent rupture of posterior follicle cells. The follicle-cell layer, devoid of the egg as a substrate, remains in the ovary and develops into a corpus luteum, similar to ovulation in mammals [24].

As mentioned previously, we validated a number of genes such as *Ance*, *Diap1*, *Ilp8*, and *Glut4EF*, which all show expression in the corpus luteum cell clusters (Fig 1E). The insulin-like peptide, *Ilp8*, involved in coordinating developmental timing, is greatly upregulated in stage 14 follicle cells and persists in corpus luteum cells [20]. The caspase binding enzyme, *Diap1*, is highly expressed in late stage (11-14) anterior follicle cells and persists in anterior corpus luteum cells [58]. The transcription factor, *Glut4EF*, shows increased expression from stage-10B main body follicle cells and reaches the highest expression level in stage-14 follicle cells and corpus luteum cells [102]. Expression of *Ance*, a gene producing an extracellular metallopeptidase, is specific to the terminal corpus luteum cells, as well as subsets of oviduct and dorsal appendage forming cells [77].

To explore cellular and transcriptomic heterogeneity of the corpus luteum, we re-clustered the 133-cell subset of corpus luteum cells from original clusters 21, 27 and 28 (Fig 6A). The cells re-clustered into 3 groups, labeled clusters 0, 1 and 2 (Fig 6B). Both *Mmp2* and *Ance* are expressed in clusters 0 and 1, indicating that they are composed of the terminal follicle cells of the corpus luteum, likely at different timepoints (Fig 7B). This also indicates that the anterior and posterior corpus luteum might be transcriptionally similar. Cluster 2 most likely represents the cells derived from main body follicle cells as they express genes such as *Ilp8* and *Glut4EF* that are expressed throughout the corpus luteum (Fig 7B). These results suggest cellular heterogeneity in the corpus luteum with specific functions of cells in different regions.

**Fig 7.**
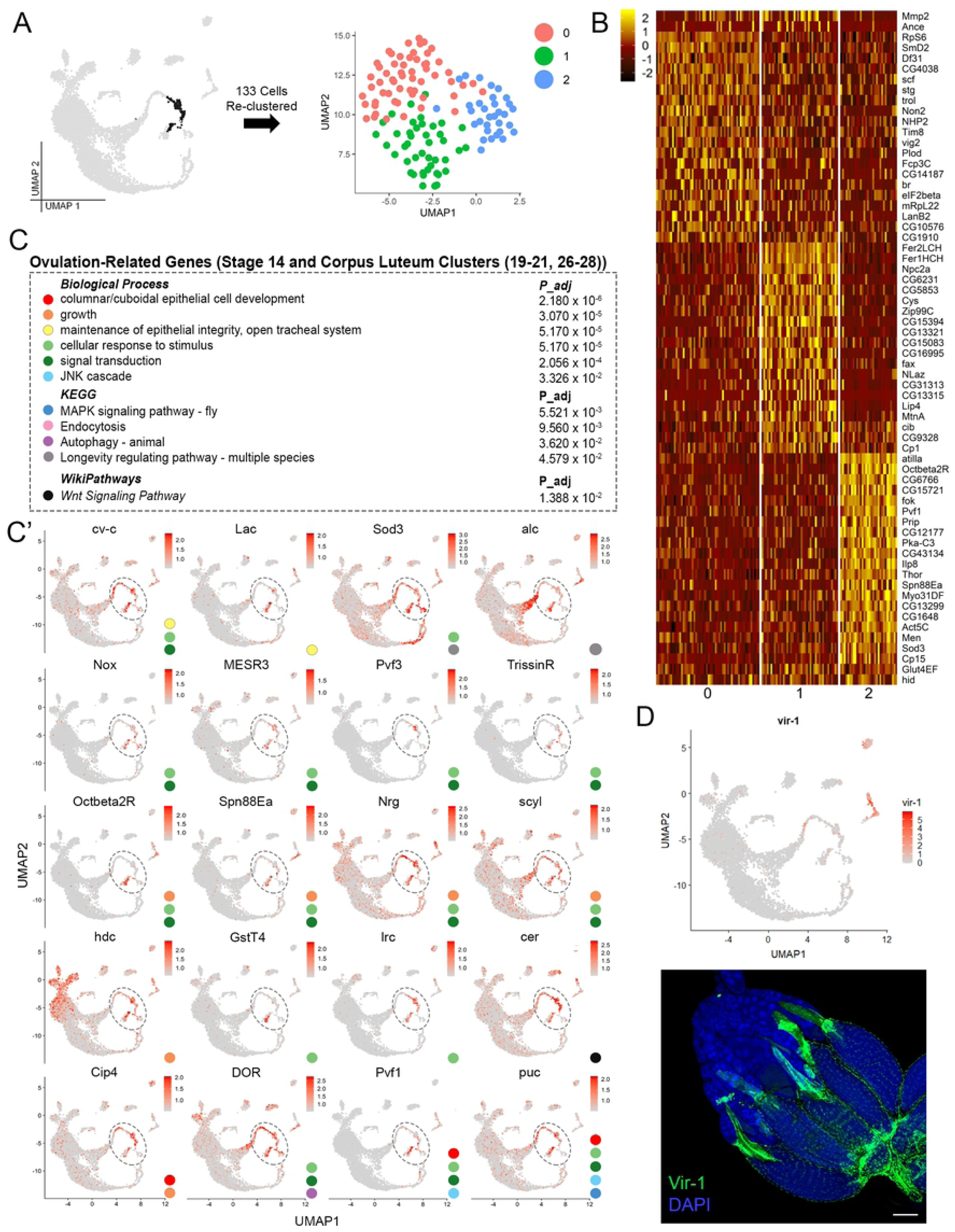
Ovulation-related genes in pre-corpus luteum cells and heterogeneity of the corpus luteum. (A) Fig 1B UMAP (grey) highlighting the 133-cell subset of corpus luteum cell clusters 21 and 27-28 (black) re-clustered at right (C) Heatmap of top 20 genes per cluster (including validated markers *Mmp2*, *Ance*, and *Glut4EF* in Fig 1) from subset plot in A. (C) GO analysis of enriched, ovulation-related genes from all stage 14 follicle cell (Stage 14 FC) also called pre-corpus luteum (pre-CL) clusters (19-20, 26) and corpus luteum (CL) clusters (21, 27-28). (C’) Feature plots of select ovulation-related genes in C. Colored circles indicate the GO term in C that each gene belongs to. Dotted ovals mark Pre-CL and CL regions of interest. (D) Experimental validation of *vir-1* (green) marked using RFP expression under T2A-Gal4 control. Expression indicated in stage 14 follicle cells before ovulation (arrow: top image) and in CL after ovulation (arrow: bottom image). Additional expression in oviduct cells indicated (*). Both images are z-projections of an entire ovary. DAPI marks nuclei. Scale bar = 100 *µ*m.

### A transcriptomic switch from oogenesis to ovulation regulation in pre-corpus luteum cells

As stated previously, corpus luteum-enriched genes, *Ilp8* and *Glut4EF*, begin their peak expression in late stage-14 follicle cells. A third, viral-response gene, *vir-1*, displays a similar pattern of sudden upregulation in stage 14 follicle cells and continued expression in corpus luteum cells after ovulation [28] (Fig 7D). Because of this shared expression timing of non-eggshell-related genes, we considered the stage-14 clusters from the stretched cell and MBFC lineage as a “pre-corpus luteum” and compared genes shared by these cells and those in the corpus luteum to gain insight into potential ovulation-related genes at the end of oogenesis.

GO term enrichment analysis of the genes identified using this method are involved in various biological processes, such as columnar/cuboidal epithelial cell development, growth, maintenance of epithelial integrity, cellular response to stimulus, signal transduction, and JNK cascade. Several key developmental pathways such as MAPK, endocytosis, autophagy, longevity, and Wnt signaling are also enriched (Fig 7C). Two of the genes identified, *Nox*, an NADPH oxidase and *Octβ2R*, an octopamine receptor, have been identified as essential for ovulation through calcium regulation in the oviduct [59, 76]. Consistent with our results, many of these ovulation-related genes also sharing expression with the cells of the oviduct and hemocyte clusters, as observed in the feature plots and *vir-1* images (Fig 7C’ and 7D).

## Discussion

In this study, we used scRNA-seq to survey the expression profiles of cells from the adult Drosophila ovary. Using a previously unreported approach, we recovered high-quality cells through removing contaminants with conflicting marker expression and experimentally validating the identity of clusters using new markers. During dissection, instead of mechanically separating intimately connected tissues (i.e. muscle sheath, hemocytes, oviduct, and fat body) from the ovary, we chose to leave them attached, including them in the dataset. Separating cells from different tissues in this way prevented damage to the ovarian cell types of interest and improved feature selection in downstream analysis. This approach allowed the clustering of all possible cell types that are physically connected to the ovary, thus taking account of cells that otherwise would have appeared as unknown contaminants. This enabled stringent fidelity assessment of cells based on an all-encompassing list of conflicting markers enabling the ultimate recovery of high-quality cells.

With a special focus on the most abundant ovarian cell type, the follicle cells, we identified their entire spatiotemporal trajectory from the stem cell niche to the corpus luteum. Using in silico subset analyses, we identified the transcriptomic basis for early differentiation of polar and stalk cells from the main body follicle cells, mitosis-to-endocycle switch, and follicular symmetry breaking. We also identified transcriptomic signatures of different follicle cell groups that carry out important developmental functions such as migration, engulfment of nurse cells, and eggshell formation. Remarkably, the dataset not only reveals a novel split in the transcriptome during symmetry breaking, but also a convergence of late-stage follicle cells as they form the corpus luteum. During this convergence, we identify ovulation-related genes in late-stage follicle cells (termed pre-corpus luteum) which may signify a novel developmental switch from oogenesis to ovulation regulation.

An unexpected advantage of this approach is the ability to analyze the relationship between ovarian and non-ovarian cell types, which show functional convergence between cells of different tissues. For example, the nurse-cell engulfing stretched cells express genes shared by the hemocytes. While some immune-related genes have been described in these “amateur” phagocytes [38], other morphology-regulating genes shared with hemocytes have not yet been identified. This introduces an interesting possibility that aspects of stretched cell and hemocyte morphology may be essential for the engulfment of cellular material, which necessitates further research. Additionally, cells in the corpus luteum possess a transcriptomic signature that has overlapping genes expressed in the oviduct cells and hemocytes, indicating a potential shared function or interaction between these cell types in regulating ovulation. This is consistent with reports in mammals that the corpus luteum functions as an endocrine body for control of reproductive timing [1, 69], and has signaling cross-talk with macrophages [14, 99]. Overall, our study provides a broad perspective of functional relatedness among cell types regulating oogenesis and ovulation. The convergence of such transcriptional “tool-kits” between developmentally unrelated cell types is an emerging theme identified using this diverse dataset. Curating information on genes that define these overlapping functions will not only help validate our current understanding of gene ontology but also identify unique genes that may have differential functions in specific cell types.

Taken together, our study provides a novel perspective of oogenesis, identifies cell-type and stage markers, and reveals functional convergence in expression between ovarian and non-ovarian cell types. Additionally, it is now possible to use this single-cell dataset to better understand the intercellular and inter-tissue signaling regulating oogenesis and ovulation.

---

**S1 Fig.**
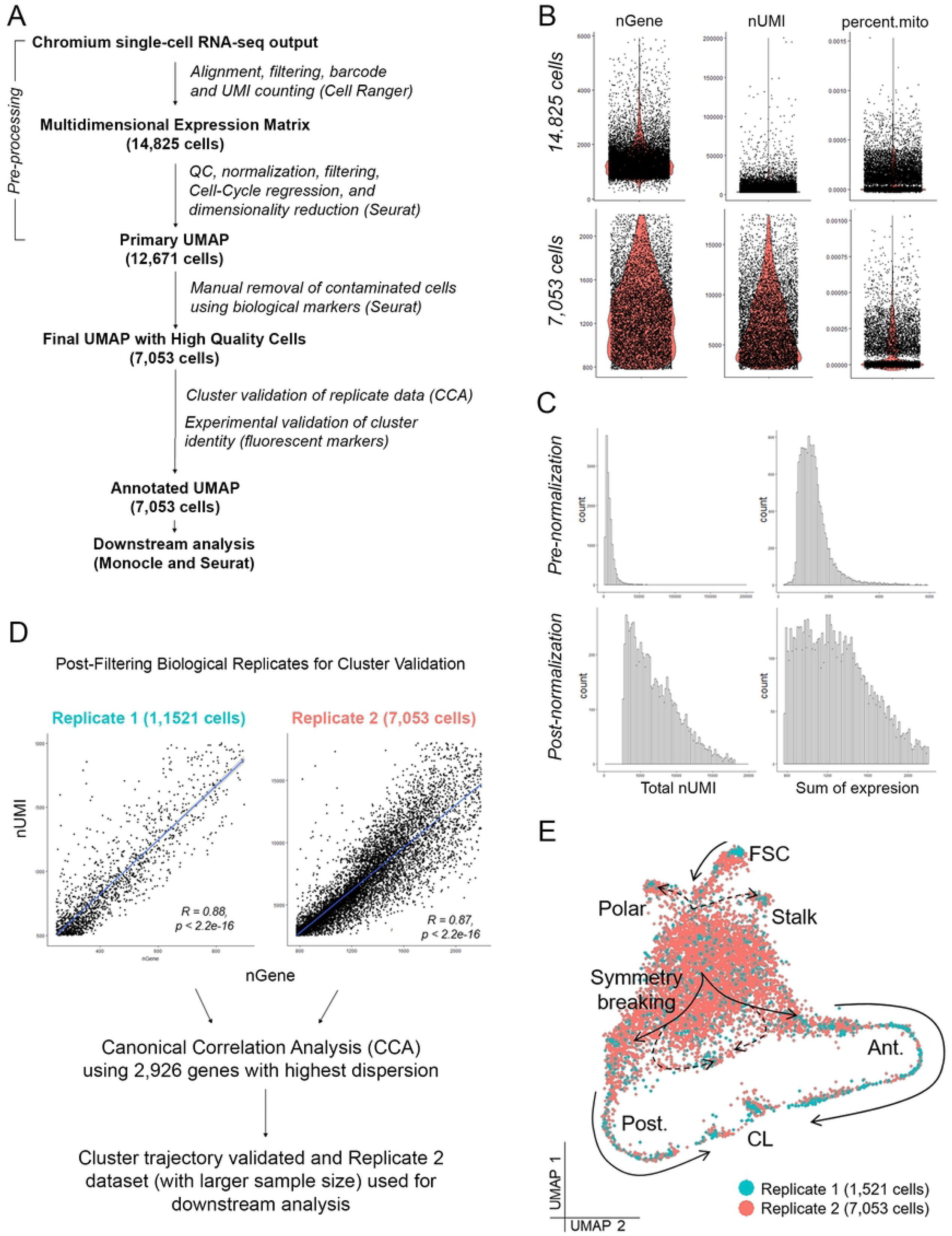
ScRNA-Seq dataset pre-processing, and verification analysis information. (A) Schematic for the scRNA-Seq analysis pipeline. (B) Violin plots for nGene, nUMI and percent.mito (percentage of mitochondrial gene expression) for dataset before (14,825 cells) and after (7,053 cells) pre-processing and manual removal of contaminated cells. (C) Feature counts were log-normalized and scaled using default options. Pre and post-normalization plots are shown for total nUMI and sum of expression. (D) Schematic used to perform Canonical Correlation Analysis (CCA) on the two biological replicate datasets: Replicate 1 (1,1521 cells) and Replicate 2 (7,053 cells). (E) UMAP of somatic/follicle cell clusters from both datasets (Replicate 1 and 2) showing the validation for fate trajectory. Cells originate from the stem cell (FSC) cluster (indicated by the solid arrow) and assume polar and stalk cell fate (indicated by the dashed arrow). The remaining cells assume mature follicle cell fate. These mature follicle cells split into two distinct transcription states (solid arrow), including various cell types in the anterior (Ant.) and posterior (Post.) during follicular symmetry breaking. Some cells from resulting anterior and posterior trajectories subsequently converge (dashed arrow) to form the migratory cells. The anterior and posterior trajectories terminally converge into the corpus luteum (CL) clusters.

**S2 Fig.**
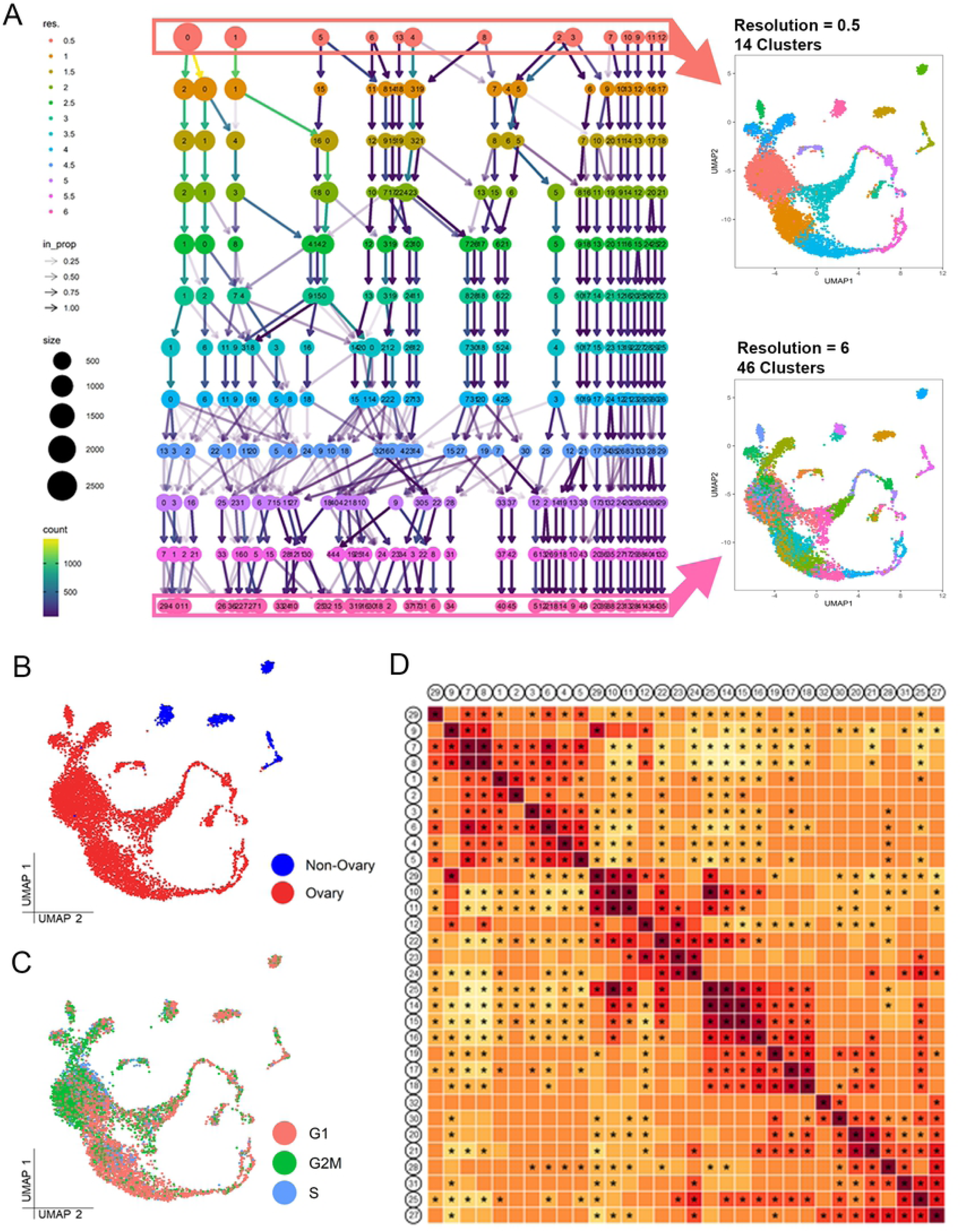
Cluster resolution and ovarian and non-ovarian cluster relationship information. (A) Clustering tree representing the relationship among all the clusters at resolutions 0.5 to 6.0. Example clustering shown for lowest (0.5) to highest (6) resolutions with cluster number ranging from 14-46. Cell type identities were resolved by separating different clusters of transcriptional states and combining the ones that had no unique markers. (B) UMAP plot showing ovarian clusters (red) including somatic and germline cell types and non-ovarian clusters (blue) including cells from oviduct, muscle, hemocytes, and fat body. (C) UMAP plot showing the cell cycle phase of all the cell clusters, based on the cell cycle score assigned for genes in S2 File. (D) Plot showing the correlation between the different cell types. Clusters are numbered according to cell type identities and numbers indicated in Figure 1B.

**S1 File. Known marker genes used to identify specific cell types and selected references where they are identified.**

**S2 File. Strategy used to separate dividing and non-dividing cells, List of genes used to assign ‘cell-cycle score’ to each individual cell.**

**S3 File. Differentially expressed genes and statistics for each cell type, as identified in Seurat (minimum expression in 25% cells of the cluster).**

## Acknowledgments

Special thanks to Roger Mercer, Yanming Yang, Cynthia Vied, Amber Brown, and Brian Washburn for their assistance in library preparation and sequencing. The authors also acknowledge Yue Julia Wang, Jerome Irianto, Michelle Arbeitman, and Jen Kennedy for help in editing and reviewing the manuscript. The authors also would like to thank Colleen Palmateer for assistance in developing the dissociation protocol and David Corcoran, Brian Oliver, Shamik Bose, Sarayu Row, Ishwaree Datta, Shangyu Gong, Chih-Hsuan Chang, and 10X support for helpful discussion, troubleshooting, inspiration, and assistance. 10X Chromium controller and other essential hardware was provided by the FSU College of Medicine Translational Science Laboratory. Special thanks to Norbert Perrimon Lab, Hugo Bellen Lab, Jin Jiang Lab and the Gene Disruption Project, who have contributed to generating transgenic lines used in our study. W.-M.D. is supported by NIH GM072562, CA224381, CA227789 and NSF IOS-155790.

## References

1. Adams EC, Hertig AT. STUDIES ON THE HUMAN CORPUS LUTEUM: I. Observations on the Ultrastructure of Development and Regression of the Luteal Cells During the Menstrual Cycle. The Journal of Cell Biology. 1969 Jun 1;41(3):696–715.

2. Arndt V, Dick N, Tawo R, Dreiseidler M, Wenzel D, Hesse M, et al. Chaperone-assisted selective autophagy is essential for muscle maintenance. Curr Biol. 2010 Jan 26;20(2):143–8.

3. Assa-Kunik E, Torres IL, Schejter ED, Johnston DS, Shilo B-Z. Drosophila follicle cells are patterned by multiple levels of Notch signaling and antagonism between the Notch and JAK/STAT pathways. Development. 2007 Mar 15;134(6):1161–9.

4. Bai H, Kang P, Tatar M. Drosophila insulin-like peptide-6 (dilp6) expression from fat body extends lifespan and represses secretion of Drosophila insulin-like peptide-2 from the brain. Aging Cell. 2012 Dec;11(6):978–85.

5. Banerjee S, Blauth K, Peters K, Rogers SL, Fanning AS, Bhat MA. Drosophila Neurexin IV Interacts with Roundabout and Is Required for Repulsive Midline Axon Guidance. J Neurosci. 2010 Apr 21;30(16):5653–67.

6. Becht E, Dutertre C-A, Kwok IWH, Ng LG, Ginhoux F, Newell EW. Evaluation of UMAP as an alternative to t-SNE for single-cell data. Bioinformatics; 2018 Apr. Available from: http://biorxiv.org/lookup/doi/10.1101/298430

7. Becker RA, Chambers author.) John, Wilks author.) Allan Reev. The new S language: a programming environment for data analysis and graphics. Boca Raton, Florida; London; New York CRC Press; 2018. Available from: https://trove.nla.gov.au/work/18397063

8. Blondel VD, Guillaume J-L, Lambiotte R, Lefebvre E. Fast unfolding of communities in large networks. J Stat Mech. 2008 Oct;2008(10):P10008.

9. Bosco G, Orr-Weaver TL. The cell cycle during oogenesis and early embryogenesis in Drosophila.. In:Advances in Developmental Biology and Biochemistry:Elsevier; 2002. p. 107–54.

10. Boyle MJ, Berg CA. Control in time and space: Tramtrack69 cooperates with Notch and Ecdysone to repress ectopic fate and shape changes during Drosophila egg chamber maturation. Development 2009 Dec 15;136(24):4187–97.

11. Burke T, Waring GL, Popodi E, Minoo P. Characterization and sequence of follicle cell genes selectively expressed during vitelline membrane formation in Drosophila. Developmental Biology. 1987 Dec 1;124(2):441–50.

12. Butler A, Hoffman P, Smibert P, Papalexi E, Satija R. Integrating single-cell transcriptomic data across different conditions, technologies, and species. Nature Biotechnology. 2018 May;36(5):411–20.

13. Buszczak M, Paterno S, Lighthouse D, Bachman J, Planck J, Owen S, et al. The Carnegie Protein Trap Library: A Versatile Tool for Drosophila Developmental Studies. Genetics. 2007 Mar;175(3):1505–31.

14. Care AS, Diener KR, Jasper MJ, Brown HM, Ingman WV, Robertson SA. Macrophages regulate corpus luteum development during embryo implantation in mice. J Clin Invest. 2013 Aug 1;123(8):3472–87.

15. Carney GE, Bowen NJ. p24 proteins, intracellular trafficking, and behavior: Drosophila melanogaster provides insights and opportunities. Biology of the Cell. 2004;96(4):271–8.

16. Casanueva MO, Ferguson EL. Germline stem cell number in the Drosophila ovary is regulated by redundant mechanisms that control Dpp signaling. Development. 2004 May 1;131(9):1881–90.

17. Cavaliere V, Donati A, Hsouna A, Hsu T, Gargiulo G. dAkt Kinase Controls Follicle Cell Size During Drosophila Oogenesis. Dev Dyn. 2005 Mar;232(3):845–54.

18. Chang Y-C, Jang AC-C, Lin C-H, Montell DJ. Castor is required for Hedgehog-dependent cell-fate specification and follicle stem cell maintenance in Drosophila oogenesis. Proc Natl Acad Sci U S A. 2013 May 7;110(19):E1734–42.

19. Cohen ED, Mariol M-C, Wallace RMH, Weyers J, Kamberov YG, Pradel J, et al. DWnt4 Regulates Cell Movement and Focal Adhesion Kinase during Drosophila Ovarian Morphogenesis. Developmental Cell. 2002 Apr 1;2(4):437–48.

20. Colombani J, Andersen DS, Léopold P. Secreted Peptide Dilp8 Coordinates Drosophila Tissue Growth with Developmental Timing. Science. 2012 May 4;336(6081):582–5.

21. D’Alterio C, Tran DDD, Yeung MWYA, Hwang MSH, Li MA, Arana CJ, et al. Drosophila melanogaster Cad99C, the orthologue of human Usher cadherin PCDH15, regulates the length of microvilli. J Cell Biol. 2005 Nov 7;171(3):549–58.

22. Dansereau DA, Lasko P. The Development of Germline Stem Cells in Drosophila. Methods Mol Biol. 2008;450:3–26.

23. Deady LD, Shen W, Mosure SA, Spradling AC, Sun J. Matrix Metalloproteinase 2 Is Required for Ovulation and Corpus Luteum Formation in Drosophila. PLOS Genetics. 2015 Feb 19;11(2):e1004989.

24. Deady LD, Li W, Sun J. The zinc-finger transcription factor Hindsight regulates ovulation competency of Drosophila follicles. Elife. 2017 19;6.

25. Deng W, Leaper K, Bownes M. A targeted gene silencing technique shows that Drosophila myosin VI is required for egg chamber and imaginal disc morphogenesis. J Cell Sci. 1999 Nov 1;112(21):3677–90.

26. Deng W-M, Althauser C, Ruohola-Baker H. Notch-Delta signaling induces a transition from mitotic cell cycle to endocycle in Drosophila follicle cells. Development. 2001 Dec 1;128(23):4737–46.

27. Diao F, Ironfield H, Luan H, Diao F, Shropshire WC, Ewer J, et al. Plug-and-Play Genetic Access to Drosophila Cell Types using Exchangeable Exon Cassettes. Cell Reports. 2015 Mar 3;10(8):1410–21.

28. Dostert C, Jouanguy E, Irving P, Troxler L, Galiana-Arnoux D, Hetru C, et al. The Jak-STAT signaling pathway is required but not sufficient for the antiviral response of drosophila. Nat Immunol. 2005 Sep;6(9):946–53.

29. Du J, Zhang J, He T, Li Y, Su Y, Tie F, et al. Stuxnet Facilitates the Degradation of Polycomb Protein during Development. Developmental Cell. 2016 Jun 20;37(6):507–19.

30. Elalayli M, Hall JD, Fakhouri M, Neiswender H, Ellison TT, Han Z, et al. Palisade is required in the Drosophila ovary for assembly and function of the protective vitelline membrane. Developmental Biology. 2008 Jul 15;319(2):359–69.

31. Eliazer S, Buszczak M. Finding a niche: studies from the Drosophila ovary. Stem Cell Res Ther. 2011 Nov 25;2(6):45.

32. Emelyanov AV, Rabbani J, Mehta M, Vershilova E, Keogh MC, Fyodorov DV. Drosophila TAP/p32 is a core histone chaperone that cooperates with NAP-1, NLP, and nucleophosmin in sperm chromatin remodeling during fertilization. Genes Dev. 2014 Sep 15;28(18):2027–40.

33. Evans CJ, Liu T, Banerjee U. DDrosophila hematopoiesis: markers and methods for molecular genetic analysis. Methods. 2014 Jun 15;68(1):242–51.

34. Gaudet P, Livstone MS, Lewis SE, Thomas PD. Phylogenetic-based propagation of functional annotations within the Gene Ontology consortium. Brief Bioinform. 2011 Sep 1;12(5):449–62.

35. Geisbrecht ER, Montell DJ. Myosin VI is required for E-cadherin-mediated border cell migration. Nat Cell Biol. 2002 Aug;4(8):616–20.

36. Glowinski C, Liu R-HS, Chen X, Darabie A, Godt D. Myosin VIIA regulates microvillus morphogenesis and interacts with cadherin Cad99C in Drosophila oogenesis. J Cell Sci. 2014 Nov 15;127(22):4821–32.

37. Grammont M, Irvine KD. Organizer activity of the polar cells during Drosophila oogenesis. Development. 2002 Nov 15;129(22):5131–40.

38. Gregory CD, Devitt A. The macrophage and the apoptotic cell: an innate immune interaction viewed simplistically?. Immunology. 2004 Sep;113(1):1–14.

39. Guruharsha KG, Rual J-F, Zhai B, Mintseris J, Vaidya P, Vaidya N, et al. A protein complex network of Drosophila melanogaster. Cell. 2011 Oct 28;147(3):690–703.

40. Hay B, Jan LY, Jan YN. A protein component of Drosophila polar granules is encoded by vasa and has extensive sequence similarity to ATP-dependent helicases. Cell. 1988 Nov 18;55(4):577–87.

41. Hill E, Broadbent ID, Chothia C, Pettitt J. Cadherin superfamily proteins in Caenorhabditis elegans and Drosophila melanogaster. Journal of Molecular Biology. 2001 Feb 2;305(5):1011–24.

42. Horne-Badovinac S. The Drosophila Egg Chamber—A New Spin on How Tissues Elongate. Integr Comp Biol. 2014 Oct;54(4):667–76.

43. Hudson AM, Petrella LN, Tanaka AJ, Cooley L. Mononuclear muscle cells in Drosophila ovaries revealed by GFP protein traps. Dev Biol. 2008 Feb 15;314(2):329–40.

44. Jang AC-C, Starz-Gaiano M, Montell DJ. Modeling Migration and Metastasis in Drosophila. J Mammary Gland Biol Neoplasia. 2007 Jun 5;12(2):103.

45. Jang AC-C, Chang Y-C, Bai J, Montell D. Border cell migration requires integration of spatial and temporal signals by the BTB protein Abrupt. Nat Cell Biol. 2009 May;11(5):569–79.

46. Jia D, Tamori Y, Pyrowolakis G, Deng W-M. Regulation of broad by the Notch pathway affects timing of follicle cell development. Dev Biol. 2014 Aug 1;392(1):52–61.

47. Jia D, Huang Y-C, Deng W-M. Analysis of Cell Cycle Switches in Drosophila Oogenesis. Methods Mol Biol. 2015;1328:207–16.

48. Johnston DS. Using mutants, knockdowns, and transgenesis to investigate gene function in Drosophila. Wiley Interdisciplinary Reviews: Developmental Biology. 2013;2(5):587–613.

49. Jordan KC, Clegg NJ, Blasi JA, Morimoto AM, Sen J, Stein D, et al. The homeobox gene mirror links EGF signalling to embryonic dorso-ventral axis formation through Notch activation. Nat Genet. 2000 Apr;24(4):429–33.

50. Kankel MW, Hurlbut GD, Upadhyay G, Yajnik V, Yedvobnick B, Artavanis-Tsakonas S. Investigating the Genetic Circuitry of Mastermind in Drosophila, a Notch Signal Effector. Genetics. 2007 Dec 1;177(4):2493–505.

51. Kim C, Han K, Kim J, Yi JS, Kim C, Yim J, et al. Femcoat, a novel eggshell protein in Drosophila: functional analysis by double stranded RNA interference. Mechanisms of Development. 2002 Jan 1;110(1):61–70.

52. Klusza S, Deng W-M. At the crossroads of differentiation and proliferation: Precise control of cell-cycle changes by multiple signaling pathways in Drosophila follicle cells. Bioessays. 2011 Feb;33(2):124–34.

53. Kolahi KS, White PF, Shreter DM, Classen A-K, Bilder D, Mofrad MRK. Quantitative analysis of epithelial morphogenesis in Drosophila oogenesis: New insights based on morphometric analysis and mechanical modeling. Developmental Biology. 2009 Jul;331(2):129–39.

54. Kongton K, McCall K, Phongdara A. Identification of gamma-interferon-inducible lysosomal thiol reductase (GILT) homologues in the fruit fly Drosophila melanogaster. Developmental Comparative Immunology. 2014 Jun 1;44(2):389–96.

55. Lantz V, Chang JS, Horabin JI, Bopp D, Schedl P. The Drosophila orb RNA-binding protein is required for the formation of the egg chamber and establishment of polarity. Genes Development. 1994 Mar 1;8(5):598–613.

56. Lasko PF, Ashburner M. The product of the Drosophila gene vasa is very similar to eukaryotic initiation factor-4A. Nature. 1988 Oct;335(6191):611–7.

57. Lee P-T, Zirin J, Kanca O, Lin W-W, Schulze KL, Li-Kroeger D, et al. A gene-specific T2A-GAL4 library for Drosophila. eLife 2018;7:e35574

58. Leulier F, Ribeiro PS, Palmer E, Tenev T, Takahashi K, Robertson D, et al. Systematic in vivo RNAi analysis of putative components of the Drosophila cell death machinery. Cell Death Differentiation. 2006 Oct;13(10):1663.

59. Lim J, Sabandal PR, Fernandez A, Sabandal JM, Lee H-G, Evans P, et al. The Octopamine Receptor Oct*β*2R Regulates Ovulation in Drosophila melanogaster. PLOS ONE. 2014 Aug 6;9(8):e104441.

60. Losick VP, Morris LX, Fox DT, Spradling A. Drosophila stem cell niches: a decade of discovery suggests a unified view of stem cell regulation. Dev Cell. 2011 Jul 19;21(1):159–71.

61. Luo L, Wang H, Fan C, Liu S, Cai Y. Wnt ligands regulate Tkv expression to constrain Dpp activity in the Drosophila ovarian stem cell niche. J Cell Biol. 2015 May 25;209(4):595–608.

62. Martin D, Zusman S, Li X, Williams EL, Khare N, DaRocha S, et al. wwing blister, a new Drosophila laminin alpha chain required for cell adhesion and migration during embryonic and imaginal development. J Cell Biol. 1999 Apr 5;145(1):191–201.

63. McKearin DM, Spradling AC. bag-of-marbles: a Drosophila gene required to initiate both male and female gametogenesis. Genes Dev. 1990 Dec;4(12B):2242–51.

64. Meehan T, F. Joudi T, Lord A, D. Taylor J, S. Habib C, Peterson J, et al. Components of the Engulfment Machinery Have Distinct Roles in Corpse Processing. PLOS ONE. 2016 Jun 27;11:e0158217.

65. Mi H, Muruganujan A, Ebert D, Huang X, Thomas PD. PANTHER version 14: more genomes, a new PANTHER GO-slim and improvements in enrichment analysis tools. Nucleic Acids Res. 2019 Jan 8;47(Database issue):D419–26.

66. Montell DJ, Rorth P, Spradling AC. slow border cells, a locus required for a developmentally regulated cell migration during oogenesis, encodes Drosophila C/EBP. Cell. 1992 Oct 2;71(1):51–62.

67. Morrison SJ, Spradling AC. Stem cells and niches: mechanisms that promote stem cell maintenance throughout life. Cell. 2008 Feb 22;132(4):598–611.

68. Murray MJ, Davidson CM, Hayward NM, Brand AH. The Fes/Fer non-receptor tyrosine kinase cooperates with Src42A to regulate dorsal closure in Drosophila. Development. 2006 Aug 15;133(16):3063–73.

69. Niswender GD, Juengel JL, Silva PJ, Rollyson MK, McIntush EW. Mechanisms Controlling the Function and Life Span of the Corpus Luteum. Physiological Reviews. 2000 Jan 1;80(1):1–29.

70. Noguerón MI, Mauzy-Melitz D, Waring GL. Drosophila dec-1 eggshell proteins are differentially distributed via a multistep extracellular processing and localization pathway. Dev Biol. 2000 Sep 15;225(2):459–70.

71. Nystul T, Spradling A. An epithelial niche in the Drosophila ovary undergoes long-range stem cell replacement. Cell Stem Cell. 2007 Sep 13;1(3):277–85.

72. Olswang-Kutz Y, Gertel Y, Benjamin S, Sela O, Pekar O, Arama E, et al. Drosophila Past1 is involved in endocytosis and is required for germline development and survival of the adult fly. Journal of Cell Science. 2009 Feb 15;122(4):471–80.

73. Park JW, Parisky K, Celotto AM, Reenan RA, Graveley BR. Identification of alternative splicing regulators by RNA interference in Drosophila. PNAS. 2004 Nov 9;101(45):15974–9.

74. Qiu X, Mao Q, Tang Y, Wang L, Chawla R, Pliner HA, et al. Reversed graph embedding resolves complex single-cell trajectories. Nature Methods. 2017 Oct;14(10):979–82.

75. Reimand J, Arak T, Adler P, Kolberg L, Reisberg S, Peterson H, et al. g:Profiler-a web server for functional interpretation of gene lists (2016 update). Nucleic Acids Res. 2016 08;44(W1):W83-89.

76. Ritsick DR, Edens WA, Finnerty V, Lambeth JD Nox regulation of smooth muscle contraction. Free Radic Biol Med. 2007 Jul 1;43(1):31–8.

77. Rylett CM, Walker MJ, Howell GJ, Shirras AD, Isaac RE. Male accessory glands of Drosophila melanogaster make a secreted angiotensin I-converting enzyme (ANCE), suggesting a role for the peptide-processing enzyme in seminal fluid. Journal of Experimental Biology. 2007 Oct 15;210(20):3601–6.

78. Rynes J, Donohoe CD, Frommolt P, Brodesser S, Jindra M, Uhlirova M. Activating Transcription Factor 3 Regulates Immune and Metabolic Homeostasis. Molecular and Cellular Biology. 2012 Oct 1;32(19):3949–62.

79. Sahu A, Ghosh R, Deshpande G, Prasad M. A Gap Junction Protein, Inx2, Modulates Calcium Flux to Specify Border Cell Fate during Drosophila oogenesis. PLOS Genetics. 2017 Jan 23;13(1):e1006542.

80. dos Santos G, Schroeder AJ, Goodman JL, Strelets VB, Crosby MA, Thurmond J, et al. FlyBase: introduction of the Drosophila melanogaster Release 6 reference genome assembly and large-scale migration of genome annotations. Nucleic Acids Res. 2015 Jan;43(Database issue):D690-697.

81. Sellin J, Schulze H, Paradis M, Gosejacob D, Papan C, Shevchenko A, et al. Characterization of Drosophila Saposin-related mutants as a model for lysosomal sphingolipid storage diseases. Disease Models Mechanisms. 2017 Jun 1;10(6):737–50.

82. Shyu L-F, Sun J, Chung H-M, Huang Y-C, Deng W-M. Notch signaling and developmental cell-cycle arrest in Drosophila polar follicle cells. Mol Biol Cell. 2009 Dec;20(24):5064–73.

83. Simões S, Denholm B, Azevedo D, Sotillos S, Martin P, Skaer H, et al. Compartmentalisation of Rho regulators directs cell invagination during tissue morphogenesis. Development. 2006 Nov 1;133(21):4257–67.

84. Stuart T, Butler A, Hoffman P, Hafemeister C, Papalexi E, Mauck WM, et al. Comprehensive Integration of Single-Cell Data. Cell. 2019 Jun;177(7):1888–1902.e21.

85. Sun J, Deng W-M. Notch-dependent downregulation of the homeodomain gene cut is required for the mitotic cycle/endocycle switch and cell differentiation in Drosophila follicle cells. Development. 2005 Oct 1;132(19):4299–308.

86. Sun J, Deng W-M. Hindsight mediates the role of notch in suppressing hedgehog signaling and cell proliferation. Dev Cell. 2007 Mar;12(3):431–42.

87. Sun J, Smith L, Armento A, Deng W-M. Regulation of the endocycle/gene amplification switch by Notch and ecdysone signaling.. J Cell Biol. 2008 Sep 8;182(5):885–96.

88. Tamori Y, Deng W-M. Tissue repair through cell competition and compensatory cellular hypertrophy in postmitotic epithelia. Dev Cell. 2013 May 28;25(4):350–63.

89. Tejedor F, Zhu XR, Kaltenbach E, Ackermann A, Baumann A, Canal I, et al. minibrain: a new protein kinase family involved in postembryonic neurogenesis in Drosophila. Neuron. 1995 Feb;14(2):287–301.

90. Timmons AK, Mondragon AA, Meehan TL, McCall K. Control of non-apoptotic nurse cell death by engulfment genes in Drosophila. Fly (Austin). 2016 Sep 29;11(2):104–11.

91. Tirosh I, Izar B, Prakadan SM, Wadsworth MH, Treacy D, Trombetta JJ, et al. Dissecting the multicellular ecosystem of metastatic melanoma by single-cell RNA-seq. Science. 2016 Apr 8;352(6282):189-96.

92. Tootle TL, Williams D, Hubb A, Frederick R, Spradling A. Drosophila Eggshell Production: Identification of New Genes and Coordination by Pxt. PLOS ONE. 2011 May 26;6(5):e19943.

93. Trapnell C, Cacchiarelli D, Grimsby J, Pokharel P, Li S, Morse M, et al. The dynamics and regulators of cell fate decisions are revealed by pseudotemporal ordering of single cells. Nat Biotechnol. 2014 Apr;32(4):381–6.

94. Tryselius Y, Hultmark D. Cysteine proteinase 1 (CP1), a cathepsin L-like enzyme expressed in the Drosophila melanogaster haemocyte cell line mbn-2. Insect Molecular Biology. 1997;6(2):173–81.

95. Tworoger M, Larkin MK, Bryant Z, Ruohola-Baker H. Mosaic analysis in the drosophila ovary reveals a common hedgehog-inducible precursor stage for stalk and polar cells. Genetics. 1999 Feb;151(2):739–48.

96. Venken KJT, Schulze KL, Haelterman NA, Pan H, He Y, Evans-Holm M, et al. MiMIC: a highly versatile transposon insertion resource for engineering Drosophila melanogaster genes. Nat Methods. 2011 Sep;8(9):737–43.

97. Witt E, Benjamin S, Svetec N, Zhao L. Testis single-cell RNA-seq reveals the dynamics of de novo gene transcription and germline mutational bias in Drosophila. Elife. 2019 Aug 16;8.

98. Wu M-F, Liao C-Y, Wang L-Y, Chang JT. The role of Slit-Robo signaling in the regulation of tissue barriers. Tissue Barriers. 2017 Jun 8;5(2).

99. Wu R, Van der Hoek KH, Ryan NK, Norman RJ, Robker RL. Macrophage contributions to ovarian function. Hum Reprod Update. 2004 Mar 1;10(2):119–33.

100. Wu X, Tanwar PS, Raftery LA. Drosophila follicle cells: morphogenesis in an eggshell. Semin Cell Dev Biol. 2008 Jun;19(3):271–82.

101. Xie T, Spradling AC. A Niche Maintaining Germ Line Stem Cells in the Drosophila Ovary. Science. 2000 Oct 13;290(5490):328–30.

102. Yazdani U, Huang Z, Terman JR. The Glucose Transporter (GLUT4) Enhancer Factor Is Required for Normal Wing Positioning in Drosophila. Genetics. 2008 Feb;178(2):919–29.

103. Zappia L, Oshlack A. Clustering trees: a visualization for evaluating clusterings at multiple resolutions. https://www.overleaf.com/project/5d93f43992c94f000143adc1 Gigascience. 2018 01;7(7).

104. Zartman JJ, Kanodia JS, Yakoby N, Schafer X, Watson C, Schlichting K, et al. Expression patterns of cadherin genes in Drosophila oogenesis. Gene Expr Patterns. 2009 Jan;9(1):31–6.

105. Zhang L, Ren F, Zhang Q, Chen Y, Wang B, Jiang J. The TEAD/TEF Family of Transcription Factor Scalloped Mediates Hippo Signaling in Organ Size Control. Developmental Cell. 2008 Mar 11;14(3):377–87.

106. Zhang Z, Stevens LM, Stein D. Sulfation of Eggshell Components by Pipe Defines Dorsal-Ventral Polarity in the Drosophila Embryo. Current Biology. 2009 Jul 28;19(14):1200–5.

